# Broad-host-range mutagenesis with CRISPR-associated transposase

**DOI:** 10.1101/2022.01.19.475551

**Authors:** Lidimarie Trujillo Rodríguez, Adam J. Ellington, Christopher R. Reisch

## Abstract

Transposons have been instrumental tools in microbiology enabling random mutagenesis, with transposons like Tn5 and Mariner, and site-specific DNA integrations with Tn7. However, programmable targeting of transposons was impossible until CRISPR-associated transposase (CasTn) systems were described. Like other CRISPR-derived systems, CasTn can be programmed with a short DNA encoded sequence that is transcribed into a guide-RNA. Here we describe a broad-host-range CasTn system and demonstrate its function in bacteria from three classes of the Proteobacteria. The CasTn genes are expressed from a broad-host-range replicative plasmid, while the guide-RNA and transposon are provided on a high-copy pUC plasmid that is suicidal in most bacteria outside of *E. coli.* Using our CasTn system, single-gene disruptions were performed with on-target efficiencies approaching 100% in the Beta- and Gammaproteobacteria, *Burkholderia thailandensis,* and *Pseudomonas putida*, respectively. The results were more modest in the Alphaproteobacterium *Agrobacterium fabrum*, with a peak efficiency of 45%, though for routine single-gene disruptions, this efficiency is adequate. In *B. thailandensis,* the system allowed simultaneous co-integration of transposons at two different target sites. The CasTn system is also capable of high-efficiency large transposon insertion totaling over 11 kbp in *P. putida*. Given the iterative capabilities and large payload size, this system will be helpful for genome engineering experiments across several fields of research.

**Significance:** The genetic modification of bacteria to disrupt native genes and integrate recombinant genes is necessary for basic and applied research. Traditional methods for targeted disruptions and insertions are often cumbersome and inefficient, limiting experiments' scale and throughput. This work developed a system for targeted transposon mutagenesis that is easy to use, iterative, and efficient. We demonstrate that the system functions across three different classes of the Proteobacteria in species widely used in research and biotechnology. Moreover, the framework of the system and accompanying plasmids that we developed will facilitate porting the system to other bacteria. Our system provides a fast and efficient protocol to genetically modify these bacteria by inserting desired genetic cargo into specific genomic targets.

## Introduction

Site-specific genome editing in bacteria has enabled research on the physiology and metabolism of bacteria while also facilitating metabolic engineering for the past three decades. Moreover, the ability to integrate genes into the chromosome has enabled heterologous expression for the production of biofuels and value-added products. In bacteria, targeted genome editing has been traditionally performed through homologous recombination (HR)^1–3^. While HR is a reliable method in some bacteria, it is often inefficient and laborious, requiring high amounts of customized editing template DNA and large numbers of cells for transformation. Construction of the editing template can be complex because long regions of homology must be cloned into a suicide vector, necessitating multi-step or multipiece assemblies^4^. In addition, multiple rounds of selection and counterselection are sometimes needed to proceed from single to double-crossover mutants.

In some bacteria, like *E. coli* and *P. aeruginosa*, there are systems for recombineering that utilize phage recombinases and enable high-efficiency integration into the genome from linear dsDNA and ssDNA^5–7^. Similarly, genome editing systems that use nucleolytic CRISPR/Cas enzymes to counter-select non-mutated cells or stimulate HR from donor DNA have been described^8,9^. While these systems are often efficient and useful, portability to other organisms can be challenging, as host-specific requirements necessitate modification of the plasmids.

Albeit not site-specific, transposons enable high throughput genomic studies and are highly useful because of the ease in identifying the insertion site^10–12^. TnSeq methods combine next-generation sequencing with transposon mutagenesis for high-throughput genome-wide studies of cell fitness, genetic interaction, and gene function^13–17^. While most transposons insert their payload DNA randomly, the Tn7 transposon behaves more like a prophage and consistently integrates into the same site. This integration is facilitated by elements TnsABCD^18^, with TnsAB acting as heteromeric transposase^19^ and TnsD mediating the site-specific recognition^20^. TnsD recognizes a conserved attachment site found across different classes of bacteria^21^ that allows insertion downstream of the gene *glmS* but is not programable to other sites^22^. While Tn7 is highly useful for efficient genomic integration of the DNA payload, the inability to program the insertion location limits its genome editing capacity. A recent phylogenetic analysis found widespread Tn7-like elements in diverse phylogenetic families, at times in association with CRISPR-Cas genes^23,24^.

Recently, two of the identified novel CRISPR-associated transposase systems were demonstrated to function as site programable transposases in *E. coli* with high on-target efficiency and large DNA payload size^25,26^. Both systems use Tn7-like elements in complex with a Cas effector. The *Scytonema hofmannii* (shCAST) system comprises a DNA targeting type V-K Cas12k in complex with the transposition proteins TnsB, TnsC, and TniQ, a homolog of TnsD. A second system, from *Vibrio cholerae*, uses a type I-F CRISPR-Cas Cascade and is similarly coupled with TniQ. Both systems resemble the Tn7 system, which includes the proteins TnsBCD^18^. The *V. cholerae* CasTn, dubbed INTEGRATE, was later shown to enable site-specific transposon integration in *Klebsiella oxytoca* and *Pseudomonas putida*^27^. For both CasTn tools, no host-specific genes are needed, potentially allowing for broader deployment than other CRISPR-enabled tools for genome editing. In addition to the efficiency of insertion, the ease with which the guide-RNA can be changed to new targets makes this system a game-changer, enabling fast and efficient mutagenesis. One caveat observed in *E. coli* was that high-overexpression of the CasTn genes increased off-target insertions. This problem is easily corrected in *E. coli* because tunable gene expression is possible by changing well-characterized promoters or altering the inducer concentration. Outside of *E. coli,* the ability to precisely control gene expression is limited, making titration of recombinant gene expression difficult^28,29^. Gene expression is also dependent on the growth state (lag, log, and stationary)^30,31^, presenting an easy way to control gene expression in addition to the promoter. Although well-characterized toolboxes exist for some organisms^32–34^, broad-host-range toolboxes show promise for identifying inducible promoters in different bacteria that can later be adopted^35^.

We sought to develop a fast, easy-to-use, and iterative broad host range system for targeted transposon mutagenesis. We chose to investigate the CasTn system from *S. hofmannii* because it has a single Cas effector protein and a relatively short functional target sequence of 24 bp coupled with an engineered single-guide RNA (sgRNA). We built several versions of the helper and donor plasmids and tested them in bacteria from three different classes of the Proteobacteria, *Agrobacterium fabrum* C58 (Af, previously A. *tumefaciens*^36^), *Burkholderia thailandensis* E264 (Bt), and *Pseudomonas putida* KT2440 (Pp). These bacteria are well-studied and relevant to biotechnology across fields like plant engineering, pathogen modeling, and bioremediation.

## Results

### Design of Broad Host-Range Plasmids and Reporter Strains

To test the *S. hofmannii* CasTn system, we first transferred the genes from the Strecker et al pHelper plasmid to a broad-host-range origin of replication. To create the CasTn plasmids (pCasTn), the P_*lac*_ promoter, *tnsB, tnsC, tniQ,* and *cas12k* genes were cloned into a plasmid with the RK2 origin of replication with the kanamycin or gentamycin resistance markers, creating plasmids pKSh2 and pKSh5, respectively. Neither pKSh plasmid nor the original pHelper possessed *lacI,* resulting in constitutive expression of the CasTn machinery from P_*lac*_. Furthermore, the plasmids designed by Leo et al. had the sgRNA encoded on the same plasmid as the CasTn genes, which prevents iterative use of the system because the plasmid cannot be easily cured after transformation. To enable an iterative mutagenesis system and facilitate easier re-targeting, we moved the sgRNA to the donor plasmid (pDonor) that possessed the transposon and replaced the R6K origin with the pUC origin of replication, which is suicidal outside of the Enterobacteriaceae. The gene encoding monomeric Red Fluorescent Protein (*mRFP*) was inserted in the place of the sgRNA target sequence. After around-the-horn cloning to add a DNA target, the *mRFP* is removed, allowing for easy identification of clones that possess re-targeted plasmids (Supplementary Figure S1). We then constructed pDonor variants with the transposon carrying tetracycline, kanamycin, gentamycin, and hygromycin markers.

### System validation by targeting *eyfp*

To enable easier testing of our system, we integrated *eyfp* into the chromosome using the site-specific mini-Tn7 integration system^37^. This allowed us to test the same target site in *Agrobacterium fabrum*, *Burkholderia thailandensis*, and *Pseudomonas putida* (Supplementary Figure S2), representing the Alpha-, Beta-, and Gammaproteobacteria, respectively. *A. fabrum* and *P. putida* were then transformed with pKSh2 and *B. thailandensis* with pKSh5 (Figure 1A). We chose targets with the KGTB PAM, because of substantial degeneracy in the first and fourth positions according to the PAM wheel^25^. The pDonor plasmid with the *eyfp-*1 sgRNA was transformed into each strain after growth to mid-log phase and spotted onto media with the corresponding antibiotic to select transposon integrants. All three transformations had about 10^3^ CFU (Colony Forming Unit) μg^−1^ of DNA transformed, indicating successful transposon integration. Mutants were assessed for loss of fluorescence on a blue light transilluminator (Figure 1B). The P_*lac*_ promoter driving transcription of *eyfp* functions across many bacterial species, though the expression is often low, and it is sometimes difficult to distinguish between cells that express *eyfp* and those that do not. Accordingly, we also screened the colonies with PCR using primers that distinguish the presence or absence of insertion based on product size (Supplementary Figure S3 A). In *B. thailandensis,* 12 of 14 colonies screened had the *eyfp* gene disrupted by the transposon (Supplementary Figure S3 B). In contrast, we could not identify any *eyfp* mutants in either *P. putida* or *A. fabrum*, suggesting that the transposon integrated at an off-target location.

**Figure 1.**
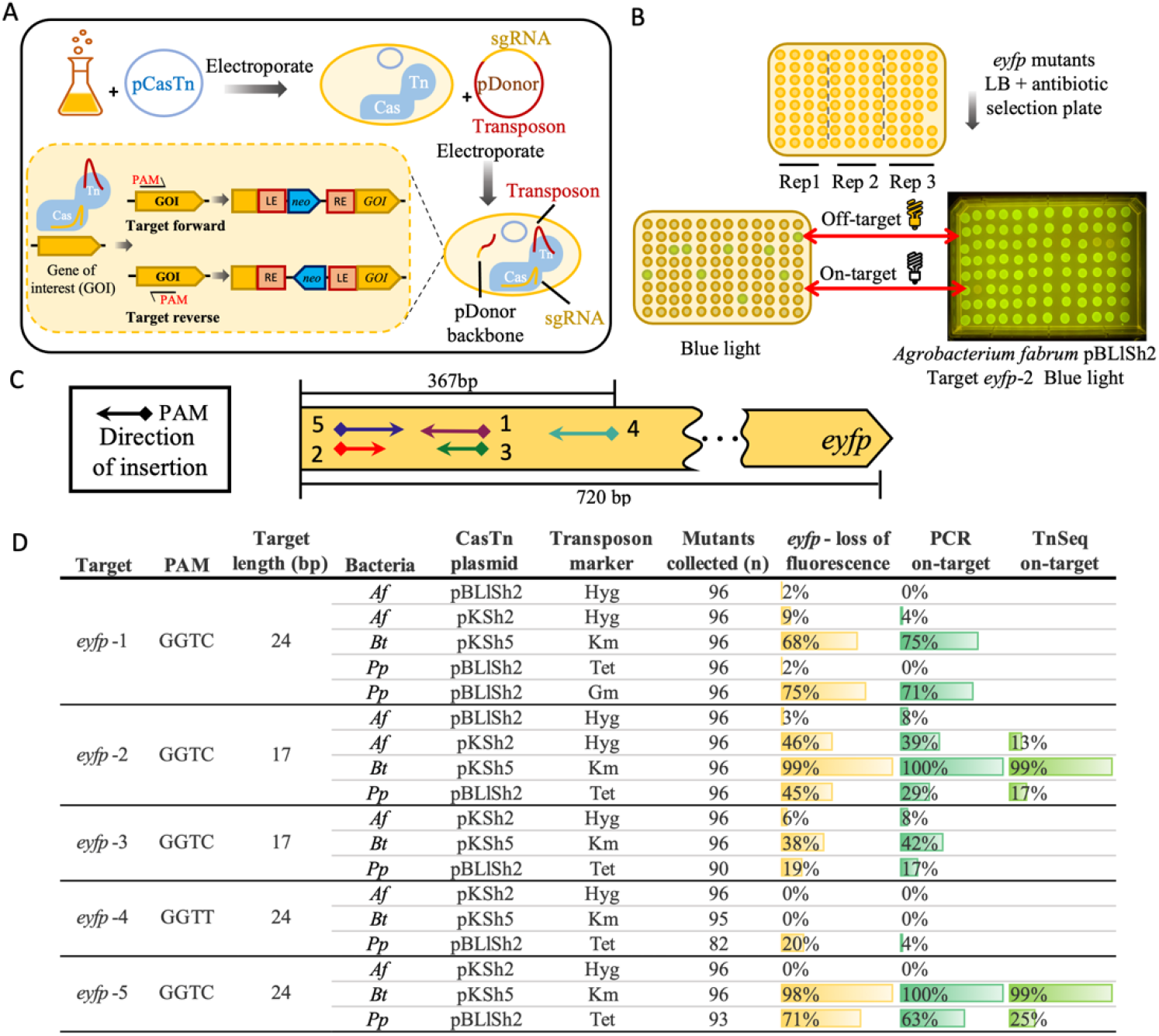
CasTn targeted disruption of *eyfp* in three Proteobacteria. **(A)** Schematic of the experimental steps for CasTn mutagenesis. The bacteria are first transformed with pCasTn, then CasTn is induced, and the strain is transformed with the suicide plasmid pDonor, which encodes for the sgRNA and the transposon. The transposon inserts directionally with the left end (LE) integrating nearest the PAM. **(B)** CasTn *eyfp* targeted integration mutants were selected by the transposon antibiotic resistance marker. The *eyfp* mutants lose fluorescence are identified upon exposure to blue light (470 nm). **(C)** Map of sgRNA targets (arrows), PAM site (diamonds), and insertion direction (arrow point). The sgRNA targeting range is above the gene and total gene length is below the gene. **(D)** Relative frequency of *eyfp* results in *A. fabrum*, *B. thailandensis,* and *P .putida*. Target name, PAM site, length in bp, the helper plasmid (CasTn plasmid), and the transposon resistance marker are given. The loss of fluorescence was calculated with n mutants collected, while genotyping was performed on 24 clones for all targets. The TnSeq on-target % was calculated with all reads from each experiment.

We hypothesized that the off-target integrations in *P. putida* and *A. fabrum* were caused by high expression of the CasTn machinery, a phenomenon reported by Strecker et al. in their description of CasTn function in *E. coli*. Accordingly, we modified the CasTn plasmid to include an inducible promoter by replacing the P_*lac*_ promoter with the P_Llac_ promoter and the *lacI* repressor, creating plasmid pBLlSh2^38^. The cells were grown overnight, subsequently diluted to an OD of 0.1, and then induced with IPTG after reaching exponential phase at an OD 0.4-0.5. After 1 hour of induction, the cells were electroporated with donor plasmids and then spotted on selective media after recovery. In addition to the original *eypf-1,* several other targets within *eyfp* were tested, including some with only 17 bp targets (Figure 1C). We chose to test different length guide-RNA targets as it has been reported that spacer length can influence the targeting in other CRISPR-Cas systems^39–44^.

In *P. putida,* some targets had over 50% efficiency (Figure 1D), while the same targets were inefficient in *A. fabrum*. For example, in *A. fabrum*, *eyfp*-1 and *eyfp*-2 yielded only 2% and 3% on-target using the pBLlSh2 CasTn plasmid, even though the same sgRNA had over 40% efficiency in *B. thailandensis* or *P. putida*, demonstrating that there is a host-specific variable affecting on-target efficiency. We then re-tested the pKSh2 plasmid in *A. fabrum* with a double setback technique, diluting to OD of 0.1 after reaching exponential phase, repeating, and then transforming with the donor plasmid during log phase. The targets *eyfp*-1 and *eyfp*-2 yielded 9% and 46% on-target insertion rates (Figure 1D), suggesting that precise conditions for growth can significantly affect the system’s functionality.

To identify the integration site of the off-targets and further confirm the on-target observations, we performed TnSeq-like mapping of the transposon and genomic DNA junction of the transformation pool. Results confirmed that in all three bacteria target *eyfp*-2, and *efyp*-5 in *P. putida* and *B. thailandensis* had high on-target insertion frequencies (Figure 1D and Supplementary Figure S4). Across all three bacteria, off-target insertions were found at low frequencies throughout their genomes. Target *eyfp-2* in *P. putida* had the highest integration at a single off-target region with 5% of total reads, compared to 17% that mapped on-target. For *B. thailandensis* and *A. fabrum,* the highest off-target regions for *eyfp*-2 account for only 0.12% and 0.31%, respectively, suggesting that off-targets do not accumulate at predetermined positions. Comparison of *eyfp*-2 on-target insertions across all three bacteria shows the peak insertion distance from the PAM for all three falls within ~20 bp of each other and spans ~300 bp for *B. thailandensis* and ~100 bp for the other organisms (Supplementary Figure S5). The differences in the insertion span length could be attributed to the difference in overall efficiency amongst the bacteria.

We also examined changing the antibiotic resistance marker on the pDonor plasmid for *P. putida*. For the *eyfp*-1 target, the on-target efficiency increased from 2% to 75% with the tetracycline marker compared to the gentamycin marker (Figure 1D and Supplementary Figure S6). Both markers resulted in similar transformation efficiencies ~10^4^ CFU μg^−1^ of DNA transformed (Supplementary Table S13), suggesting that selection strength alone was not the reason for the observed discrepancy. The tetracycline marker was efficient in other instances, discussed below with amino acid biosynthesis gene targets.

Overall, the most efficient system in each bacteria had an average transformation efficiency of 10^4^ CFU μg^−1^ DNA, allowing the retrieval of thousands of transposition mutants with just one mL of cell culture (Supplementary Table S8). There was no significant difference regarding the on-target efficiency of the different length sgRNA that we tested (p-value > 0.05, Kruskal-Wallis test; Supplementary Table S9). Moreover, the different targets had varying on-target efficiency in all the bacteria tested. Overall, results had an agreement of 94% for the paired phenotypic and genotypic tests confirming or refuting on-target integration. Differences between loss of fluorescence (n=96) and PCR (n=24) percentages may be due to the difference in sample size, but these present with strong positive correlation across all bacteria (r = 0.9823, p-value = < 2×10^−16^; Supplementary Table S10). Because each bacterium showed between 40-99% efficiency for at least one sgRNA target, we were encouraged to extend this system to different targets. In practical terms, efficiencies of even 40% allow for rapid, facile screening of mutants, as screening only three colonies will enable the identification of a mutant with the correct insertion.

### Testing *B. thailandensis* with auxotrophic targets

We next sought to confirm that the CasTn system could disrupt genes with physiological relevance. Accordingly, we targeted genes whose disruption would cause amino acid auxotrophy. Disruption of these genes would provide two lines of evidence that the integrations were on-target since mutants would be unable to grow on a minimal medium but have growth restored with the amendment of the cognate amino acid (Figure 2A and B). To minimize the chance of pleiotropic effects changing the cell physiology, we targeted genes on single-transcriptional units or last in the predicted operon.

**Figure 2.**
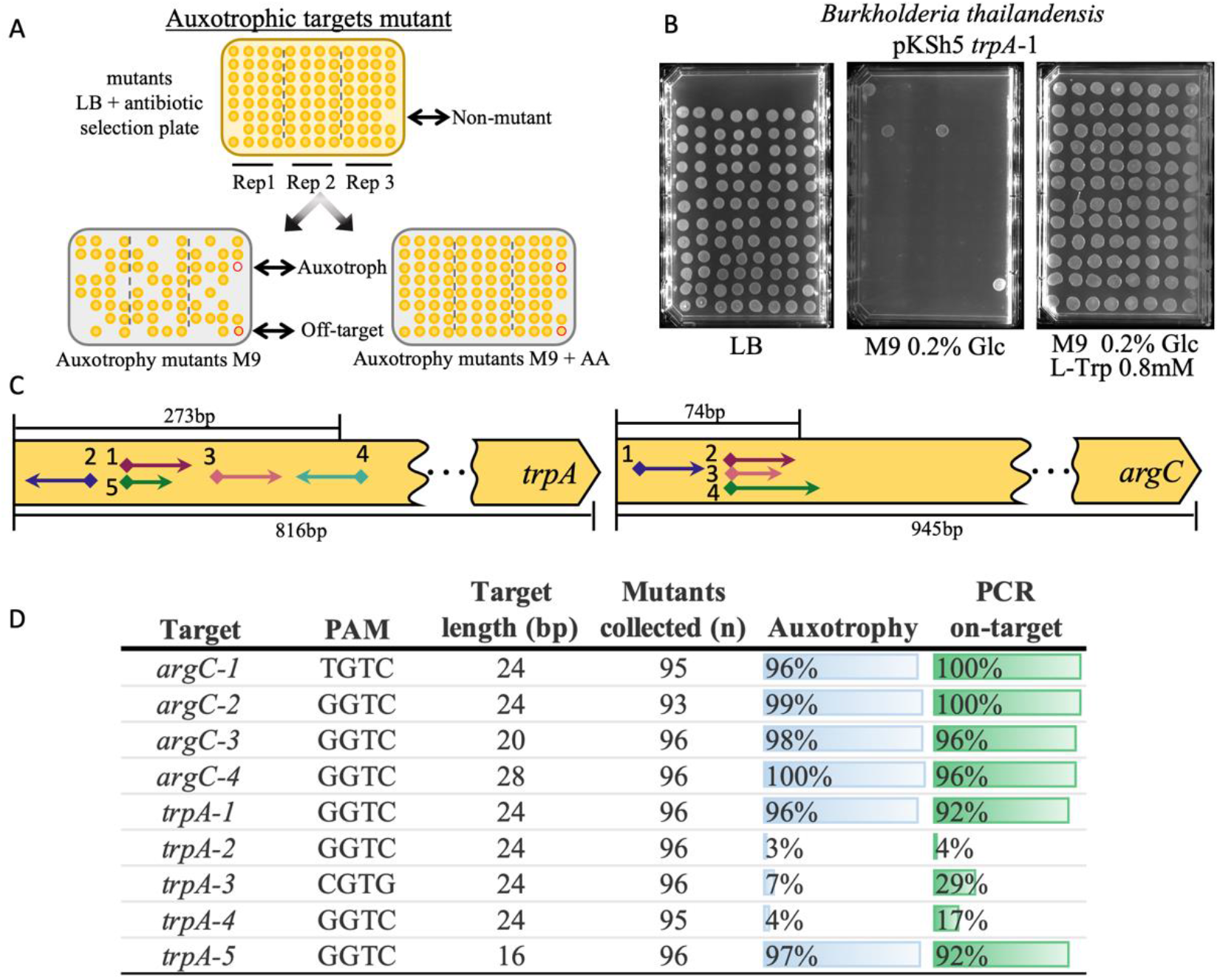
Auxotrophic screening and target efficiency in *B. thailandensis*. **(A)** Auxotrophic CasTn mutants with the integrated resistance marker are spotted onto LB agar plate plus antibiotic. Subsequently, mutants are passed to M9 agar with and without the cognate amino acid. (**B)** Selection plates for *B. thailandensis trpA*-1 target that results in tryptophan auxotrophy. M9 minimal media plates with 0.2% glucose (Glc) with or without 0.8 mM tryptophan (L-Trp). **(C)** Map of sgRNA targets (arrows), PAM site (diamonds), and insertion direction (arrow point). (**D)** Relative frequency table of on-target gene disruption results with auxotrophy complementation (n= mutants collected) and genotype (n=24).

In *B. thailandensis,* we targeted *argC* and *trpA* to yield arginine and tryptophan auxotrophs respectively. A total of five *trpA* and four *argC* targets located within the first 33% of the coding sequence were tested (Figure 2C). Delivery of the pDonor plasmids resulted in an average transformation efficiency of 8.3× 10^3^ CFU μg^−1^ across all targets (Supplementary Table S8). The colonies were then resuspended in LB and spotted onto LB agar, incubated overnight, and then spotted again on M9 medium with glucose and with or without the auxotrophic amino acid (Figure 2B). The data summarized in Figure 2D shows that six of nine sgRNA’s had very high efficiency with over 90% on-target. Both *trpA*-1 and *trpA-*5, which targeted the same region but with a 24- or 16-bp protospacer, yielded very high on-target efficiencies, confirming that protospacers of just 16-bp are functional. The other three *trpA* sgRNA, each with a 24-bp protospacer and targeting regions slightly up- and downstream, had much lower efficiencies.

All four targets in *argC* yielded very high on-target efficiencies. We observed that the rich medium did not provide sufficient arginine during preliminary experiments, and the auxotrophs grew poorly. Thus, in subsequent experiments, arginine was added to the recovery medium. Target *argC*-1, which had a TGTC PAM site and 24-bp protospacer, was nearly 100% efficient. The *argC*-2 target had a GGTC PAM, and a 24-bp protospacer was similarly efficient. We then tested the same PAM as the *argC*-2 region, but with different length sgRNA, *argC*-3 (20-bp) and *argC*-4 (28-bp), and in both cases found nearly 100% on-target efficiency on both tests.

In most cases, the PCR confirmed the observed phenotype with 96% of PCRs in agreement with the auxotrophy assessment and with a positive Pearson correlation of r = 0.9875, p-value = 1.25×10^−12^ (Supplementary Table S10). Two factors likely caused the slight differences between phenotype and genotype efficiencies for targets trpA-3-4. First, there was variation in sample size since only a quarter of the mutants screened for auxotrophy were also genotyped. Second, there appeared to be some mutants that were mixed populations that had both auxotrophs and prototrophs. The PCR indicated a mutant genotype with these samples, though the putative mutant also grew on M9.

### Testing *P. putida* with auxotrophic targets

In *P. putida,* we targeted *trpF* and *serA* to make tryptophan and serine auxotrophs, respectively (Figure 3A). Mutants obtained with target *serA*-1 had high rates of auxotrophy, while targets *serA*-2 and *serA-*3 were less than 10% on-target (Figure 3B). The TnSeq results also showed low efficiency for *serA-*2, with on-target reads accounting for only 0.12% of total mapped reads and infrequent transposon insertion around the on-target area (Supplementary Figure S7A and B). For *serA-*2 (24-bp target), we then tested versions of the guide that were 20- and 28-bp in length, with *serA*-4 and *serA-*5, respectively. Comparison of the three sgRNA showed virtually no difference in the number of auxotrophic mutants obtained from each (Figure 3B), suggesting that adjusting sgRNA length does not improve on-target efficiency in *P. putida*.

**Figure 3.**
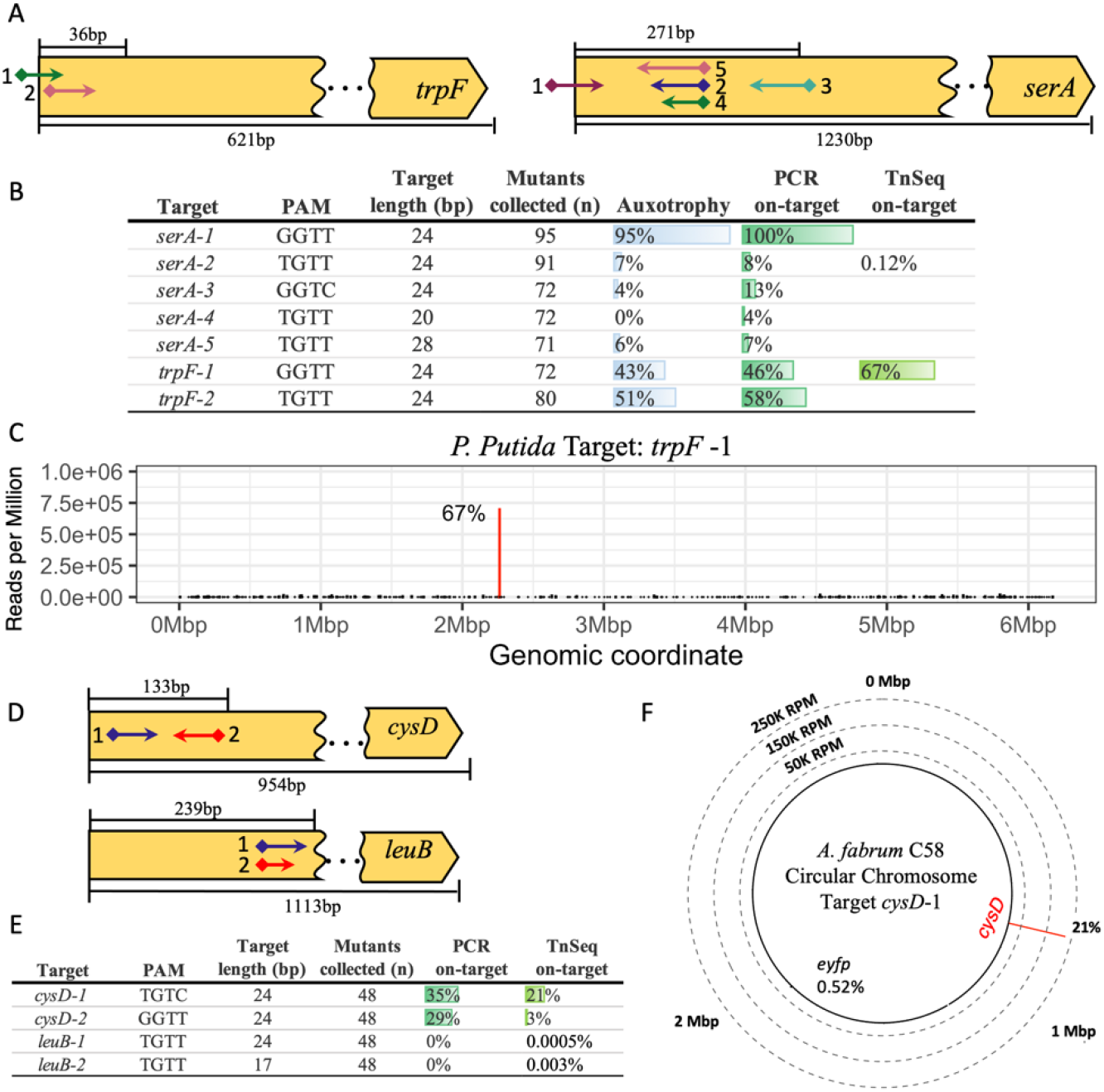
Gene disruption in *P. putida* and *A. fabrum*. **(A)** Map of auxotrophic sgRNA targets across *P. putida* amino acid biosynthesis genes *trpF* and *serA*. SgRNA targets (arrows) with the PAM site (diamonds) and insertion direction (arrow point). SgRNA targeting range (above gene) and total gene length (below gene). **(B)** Relative frequency table of auxotrophy complementation (n= mutants collected), genotype (n=24) and TnSeq results for *P. putida* targets. **(C)** TnSeq results for *P. putida* target *trpF*-1 with on-target reads (red bar) and frequency (%) across the whole genome. **(D)** Map of sgRNA targets across *A. fabrum* amino acid biosynthesis genes *cysD* and *leuB*. **(E)** Relative frequency table of genotype (n=48) and TnSeq results for *A. fabrum* targets. **(F)** TnSeq results for *A. fabrum* target *cysD*-1 with on-target reads (red bar) and frequency (%) across the whole genome. Dashed rings represent the Y-axis in reads per million (RPM) with Mbp (Megabase pair) denoting the chromosome coordinates.

In contrast, both *trpF* targets had a moderate frequency of auxotrophy. The *trpF*-1 had 43%, and *trpF*-2 had 51% auxotrophy, consistent with the genotyped on-target frequencies (Figure 3B). The TnSeq data for target *trpF*-1 shows 67% on-target reads with insertions spanning ~100 bp of the target site (Figure 3C and Supplementary Figure S7C). The TnSeq data revealed no prominent off-target locations for target *trpF*-1; instead, off-targets were spread throughout the genome. For *trpF-*1 and *serA-*2, the highest off-target sites found through TnSeq account for 1.73% and 3.76% of all insertions. Overall, the phenotype and genotype results for *P. putida* had good agreement, with 91% of screened clones having matching phenotype and genotype, with a Pearson correlation of r = 0.9734, p-value = 1.06×10^−09^ (Supplementary Table S10).

### Targeting amino acid biosynthesis genes *A. fabrum*

In *A. fabrum,* the cysteine and leucine biosynthesis genes, *cysD* and *leuB* (Figure 3D), were selected as possible auxotrophic targets. It was previously found that *cysD* disruption resulted in mucoid colonies but not auxotrophy on agar plates^45^. However, this phenotype was challenging to discern, and *leuB* disruptions were also prototrophic under the conditions tested here. Thus, genotyping by PCR was used to quantify transposons insertions (Figure 3E). The average transformation efficiency for these targets ranged from 10^3^-10^4^ CFU μg^−1^ of DNA transformed (Supplementary Table S11).

Genotyping revealed that targets in *cysD* had transposon insertion frequencies over 25% for both *cysD*-1 and *cysD*-2. TnSeq data showed a similar result for *cysD*-1, with an on-target insertion frequency of 21% (Figure 3F, Supplementary Figure S8). However, *cysD*-2 had only 3% on-target based on the TnSeq data (Figure 3E), though this was still the most abundant integration site (Supplementary Figure S9). This slight discrepancy is likely attributed to amplification bias during the Tn-Seq mapping procedure^46^. For targets *leuB*-1 (24-bp) and its shorter version *leuB*-2 (17-bp), no on-target transposon insertions were found through genotyping by PCR (n=48), and TnSeq found <0.005% integrations in the targeted region of *leuB* for both targets (Figure 3E and Supplementary Figure S10B). The *A. fabrum* TnSeq results had similar off-target patterns for all four sgRNA, with integrations mapping across the whole genome, encompassing both the circular and linear chromosomes and the native pAT and pTi plasmids. The most prominent location for off-target integration for all four sgRNA was the *eyfp* integration region, though only <1% of reads were mapped to that location for all targets (Figure 3F and Supplementary Figures 8-10A).

The relatively high level of on-target integrations observed for *cysD* are promising, though improvements are needed to enable a consistently efficient system. Perhaps further optimization of the promoter driving the CasTn expression or growth conditions would further improve efficiency. Alternatively, further experimentation could identify properties of targets or PAM’s most likely to function with high efficiency using the plasmids developed here.

### Multiplex targeting in *B. thailandensis*

Given the high efficiency of insertion in *B. thailandensis,* we investigated whether simultaneous transposon insertion at two different target sites was possible. Plasmids targeting *efyp*-2 and *trpA*-1 with kanamycin and tetracycline resistance markers were co-transformed into cells that possessed the pKSh5 plasmid. Colonies appeared two days after plating on LB with selection for both resistance markers, with a transformation efficiency of 1.2×10^5^ CFU μg^−1^ DNA transformed (Supplementary Table S12). Colonies were spotted onto LB and minimal medium to observe the phenotype, and remarkably, most colonies had both mutations (Figure 4A). Transposon sequencing of the transformed pool confirms this trend (Figure 4A and B), where both mutations account for a total of 99% of reads in the dataset across both chromosomes. The highest off-target region accounts for only 0.61% of all reads mapped across the genome. We attribute the slight unevenness of the on-target numbers, 43% and 56% of reads for *eyfp* and *trpA,* respectively, to bias in the TnSeq amplification step. Nonetheless, the ability to introduce two insertions simultaneously will streamline strain engineering applications and profoundly increase the ease of strain construction.

**Figure 4.**
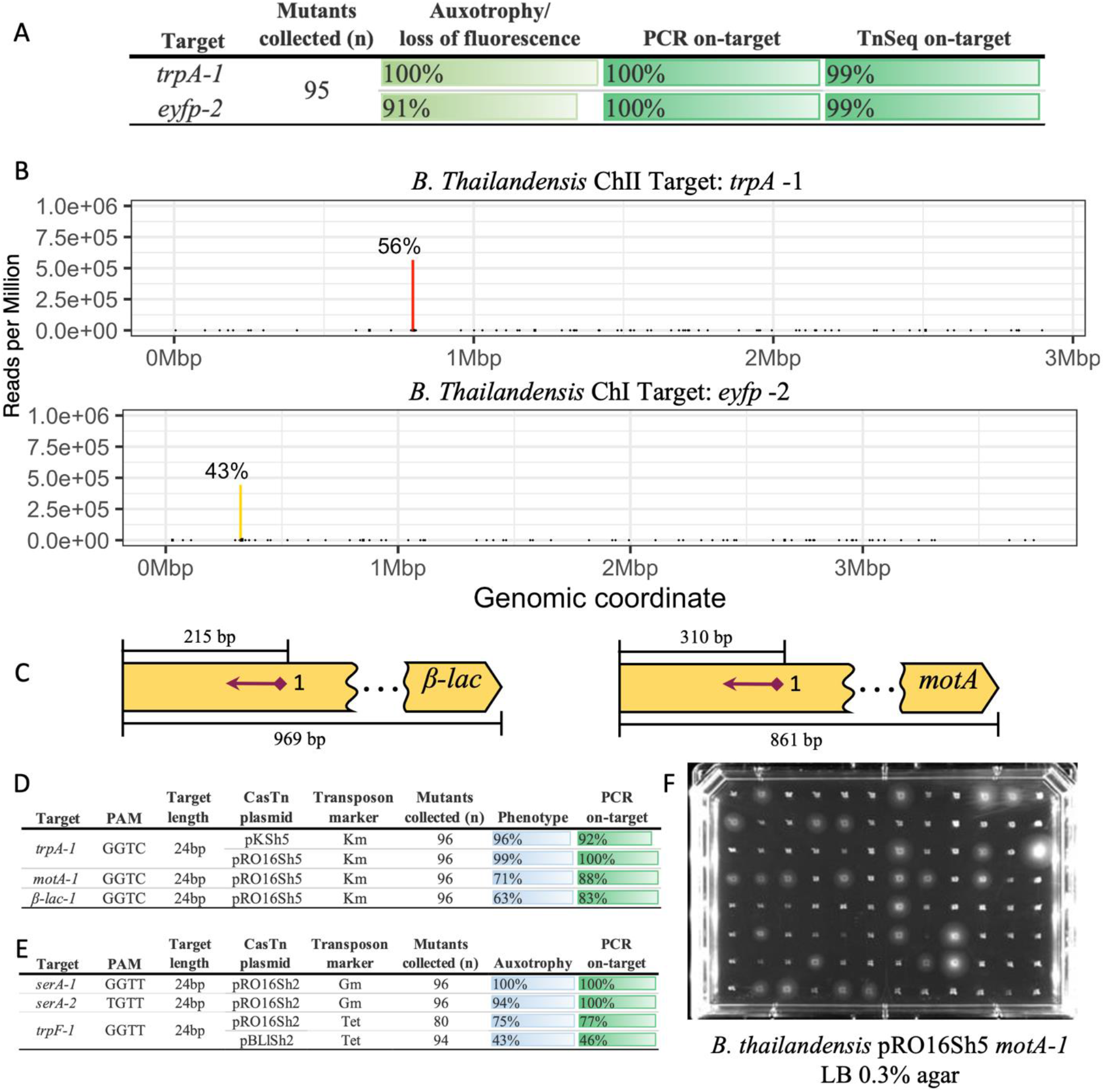
Multiplex Tn Integrations in *B. thailandensis* and temperature-sensitive pCasTn. **(A)** Relative frequency table of multiplex targeting with auxotrophy/fluorescence disruption and genotyping results for *B. thailandensis* in co-transformed targets *trpA*-1 and *eyfp*-2. **(B)** TnSeq results for *eyfp*-2 (yellow) in chromosome I and *trpA*-1 (red) in chromosome II. Read frequency presented as a percentage across both chromosomes. **(C)** Map of sgRNA targets in *B. thailandensis* genes *motA* and class A β-lactamase (β-lac) encoding BTH_RS07435. **(D)** Relative frequency table for *B. thailandensis* pRO16Sh5 temperature-sensitive pCasTn with phenotype (n= mutants collected) and genotype (n=24) results. Phenotype results reflect disruption of tryptophan production (*trpA-1*), motility (*motA-1*) or carbenicillin 100 μg/ml resistance (class A β-lactamase). **(E)** Relative frequency table for *P. putida* pRO16Sh2 temperature-sensitive pCasTn with auxotrophy (n= mutants collected) and genotype (n=24) results. **(F)** Motility assay for pRO16Sh5 *B. thailandensis motA-1* in semi-solid LB 0.3% agar, successful gene disruption results in loss of motility halo.

### Assessment of whole plasmid integration

It was observed that the ShCAST sometimes resulted in co-integrates, where the entire donor plasmid is integrated into the target site because the system lacks an equivalent of the endonuclease *tnsA*^19,47–49^. To determine the frequency of co-integrate formation, we performed PCR with primers that amplify the plasmid backbone outside of the transposon(Supplementary Figure S11A). Consistent with the observation in *E. coli*, the plasmid was sometimes integrated into the chromosome in all three strains. In *B. thailandensis* and *P. putida*, 43% and 32% of the integrations possessed the whole plasmid, respectively. In comparison, *A. fabrum* had the most plasmid integrations, with an average of 81% (Supplementary Table S14 and Supplementary Figure S11B).

Although integration of the whole plasmid is not desired, disruption of the targeted genes occurs nonetheless. Thus, this phenomenon does not detract from the usefulness of this targeted mutagenesis system. Since the antibiotic marker is flanked with FRT recognition sites, we suspected that the flippase (Flp) recombinase would remove the plasmid backbone and marker simultaneously, resulting in the desired genotype. We first confirmed that Flp recombinase worked as expected for the excision of the selection marker from a transposon integration in *B. thailandensis* without the plasmid backbone (Supplementary Figure S12). We next tested whether Flp recombinase could remove both integrated markers and the plasmid integration. PCR amplification of a co-integrate confirmed that about two-thirds of colonies were kanamycin-sensitive and had lost the plasmid backbone (Supplementary Figure S11C and D). Those colonies that still had resistance after induction of Flp were found to have retained one of the Kanamycin markers copies while the other had been flipped; other times, the pUC backbone was also found through PCR, possibly from inefficient excisions (Supplementary Figure S11E). Nonetheless, these results indicated that successful marker excision could remove the pDonor backbone in cases of complete plasmid integration with high efficiency.

### Insertion of an 11 kbp operon in *P. putida*

In *E. coli* payloads of 10 kbp were efficiently integrated into the genome^27^. To assess whether these large insertions were also possible in *P. putida*, we tested a transposon that totaled 11.1 kbp containing a gentamycin resistance gene and the violacein synthesis pathway composed of *vio*ABCDE, which produces a purple pigment in cells (Supplementary Figure S13A). This transposon was delivered from a pDonor containing a sgRNA targeting *eyfp*-1, with the complete vector totaling 13.9 kbp. We successfully transformed and recovered violacein transposon mutants, with a mean transformation efficiency of 7.7×10^3^ CFU μg^−1^ DNA transformed (Supplementary Table S13). All mutants presented with violet color at the time of recovery, with 91.5% of all mutants collected (n=47) found to be on-target when genotyped (Supplementary Figure S13B). We attempted the same experiment in *B. thailandensis* but were unsuccessful, though we were also unable to transform a replicative plasmid with *vio*ABCDE.

### Temperature-sensitive pCasTn

Our goal was to improve the mutant construction process and allow easy curing of the CasTn plasmid with a temperature-sensitive origin of replication (Ts). Accordingly, we moved the pCasTn to the pRO1600 Ts origin previously used in *B. thailandensis*^50^. The resulting plasmids, pRO16Sh5 (Gentamicin) and pRO16Sh2 (Kanamycin) were transformed into *B. thailandensis* and *P. putida*, respectively. For the non Ts plasmids in these two organisms, curing pCasTn took three days in *B. thailandensis* and *P. putida* and resulted in ~70% of the population losing the plasmid (Supplementary Figure S14A and C). In *A. fabrum,* increasing the incubation to two overnights resulted in a significant increase in pCasTn loss from <10% to ~75% of the population (Supplementary Figure S14D). With pRO16Sh5, four hours of growth at a high temperature (42°C for Bt and 37°C for Pp) resulted in over 85% plasmid loss in both bacteria (Supplementary Figure S14B and E), thereby decreasing the time needed for curing from three days to one day. The genome integration efficiency using pRO16Sh5 was assessed in *B. thailandensis* using the target *trpA-1* and two new physiologically relevant targets for motility (*motA-1*) and β-lactam resistance (β-lac-1) (Figure 4C). Disruption of *motA-1* causes loss of the motility halo in semi-solid medium (Figure 4F) while target β-lac-1, which encodes for a class A β-lactamase, renders carbenicillin sensitivity. The *trpA-*1 target had similar results with the pKSh5 and pRO16Sh5 plasmids, with an on-target efficiency of ~100% (Figure 4D). The pRO16Sh5 system also had high efficiency for targets *motA-1* and β-lac-1, with over 60% on-target efficiency in both (Figure 4D). In *P. putida* the on-target efficiency with pRO16Sh2 was also high (Figure 4E). In comparison to the pBLlSh2 plasmid using a tetracycline marker, an improvement from ~45% to ~75% efficiency with *trpF*-1 was observed. We then tested the *serA*-1 and *serA*-2 targets with the gentamicin marked transposon. Both targets had efficiencies near 100% with the pRO16Sh2 plasmid, a dramatic increase from the <10% efficiency for target *serA*-2 with a tetracycline marker in the pBLlSh2 system. Though we mostly attribute this increase to the gentamicin marker since a similar increase in efficiency was observed relative to the tetracycline marker for target *eyfp*-1. Nonetheless, this data demonstrates that the pRO16Sh plasmid allows for a simplified mutagenesis protocol with on-target efficiency meeting or exceeding the pBLlSh2 plasmid.

## Discussion

Random transposon mutagenesis has allowed the study of gene disruptions that cause loss of function in bacterial cells. The randomness of integration has served as a valuable tool to rapidly create whole-genome deletion libraries that can be screened for loss of function or population-level dropout analysis^51–53^. Until recently, this technology was limited by the inability to target specific locations, making retrieval of specific mutants a lengthy and challenging process as transposon libraries had to be created and mapped to find desired mutants. CasTn technology combines the efficiency of transposon mutagenesis with the site-specific targeting of CRISPR. Our approach uses CasTn from *S. hofmannii* for an iterative and easily re-programmable transposon mutagenesis system. Re-targeting the pDonor sgRNA through around-the-horn cloning is an easy and efficient two-step process of amplification and ligation that can be parallelized for high-throughput plasmid construction. Furthermore, the Ts CasTn plasmids in *B. thailandensis and P.* putida maintain high efficiency and streamline the plasmid curing process. Although this work focused on only three members of the Proteobacteria, we expect that the system and plasmids will be useful in additional bacteria.

Overall, our approach allowed targeted transposon mutants in a short time, with initial set-up and one round of integration taking as little as three days. While the on-target efficiencies we observed varied widely, the ease with which target plasmids can be constructed, the efficiency of transformation, and the large payload size make this technique potentially transformative. We demonstrated that on-target efficiency could be improved by changing promoters that drive the expression of the CasTn genes. We found multiple targets with high ~100% on-targeting efficiency in *B. thailandensis* and *P. putida* and over 40% efficiency for *A. fabrum*. These efficiencies allow quick identification of mutants because even 10% efficiency requires only ten colonies be screened to identify one with the desired mutation. We also obtained multiplex transposition into two separate target sites with different resistance genes, demonstrating the capacity to achieve multiple targeted genomic modifications in tandem in a cell, considerably reducing the time needed to create polygenic mutants. Finally, curing the CasTn plasmid for a finished mutant in all three bacteria can be achieved in as little as one day with the use of the Ts pRO16Sh5 in *B. thailandensis* and *P. putida* curing ~100% of the population, or in three to four days with the other CasTn plasmids resulting in ~70% efficiency.

There are several advantages of the CasTn system compared to other methods for genome editing in bacteria. Compared to traditional homologous recombination (HR) from suicide vectors, the CasTn protocol is faster and less laborious because custom vectors with long regions of homology do not need to be constructed (Supplementary Figure S16). Instead, only 16-24 bp of the sgRNA on pDonor is changed, a modification that is both easy to perform and scalable for high-throughput mutagenesis.

Additionally, the transformation efficiency allows the scale to be decreased by order of magnitude compared to other engineering methods. HR protocols often require 20-50 mL of culture concentrated 200x for transformation or large concentration DNA (>500ng) to achieve high efficiencies^54–56^. We used less than 1 mL of cell culture for the three bacteria tested here and with as little as 50 ng of DNA to obtain up to 10^5^ CFU’s. These decreased volumes and concentrations allow for simpler workflows and easy scaling to test many sgRNA or target many genes.

Methods for recombineering with phage-derived proteins have been described in all three bacteria tested here, although in *A. fabrum*^57^ these methods have not been widely adopted. The system described in *B. thailandensis*^58^ still requires a complicated cloning step to integrate regions of homology and multiple rounds of selection/counterselection. In *P. putida*, several recombineering techniques have been described^56,59^, some of which use CRISPR/Cas to counterselect^60^ against unedited cells or stimulate recombination^32,61^. These techniques are highly useful, and some enable scar-free mutagenesis. However, each method has its downsides, including low editing efficiency, the limited size of integrations, and complicated cloning schemes. While the CasTn also has downsides, the large cargo size and relative simplicity in targeting are inherent advantages. In contrast to other CRISPR-enabled methods that require the design and construction of both the guide-RNA and the cognate repair template, the CasTn system requires customization of only the guide-RNA. Overall, compared to HR and recombineering, while CasTn does not currently support scarless mutations, it simplifies and accelerates multiplex genomic modifications while maintaining site-directed mutagenesis.

In comparison to other CasTn systems (Supplementary Figure S17), the type I-F *V. cholerae* (Tn6677), INTEGRATE (VchINT)^27^, was recently demonstrated to function from a broad host range vector and was highly efficient in *Klebsiella oxytoca* and *P. putida.* This system used a replicative plasmid encoding the CasTn, sgRNA, and the transposon (pSPIN) or a two-plasmid system with the transposon encoded in a second plasmid. Because the pSPIN plasmid is replicative, it is unclear if the plasmid can be easily cured in the cells which have integrated the transposon. Without curing this plasmid, only one round of mutagenesis can occur. In contrast, our two-plasmid system has a replicative plasmid carrying the CasTn machinery, while the sgRNA and transposon are encoded on a suicide vector. This design strategy makes our system inherently iterative because non-integrated donor DNA cannot be maintained, enabling faster subsequent mutagenesis with a compatible antibiotic marker and different sgRNA targets. Consistent with previous observations, we also found that the *S. hofmannii* CasTn inserts transposons directionally with the left arm adjacent to the target PAM. On the other hand, the *V. cholerae* derived system inserts the transposon in both orientations, making confirmation of on-target insertion more difficult.

The CasTn plasmids described here, available with several antibiotic resistance markers, can potentially be used directly in other members of the Proteobacteria. As we observed, initial optimization of growth and target selection may be necessary to find conditions suitable for high on-target transposition in other organisms. To this end, our recently described plasmid toolbox^35^ can be used to identify regulated promoters that function in the Proteobacteria to match the level of expression required for high on-target efficiency. Further research and more comprehensive studies on the target efficiency across the genome could improve the ability to predict which target sites will function with higher efficiency. In sum, the CasTn system described here is a fast, easy, and efficient solution to creating gene disruptions and insertions with a wide range of applications suitable for deployment across diverse bacterial species. While initial work is needed to set up the system, the potential to modify non-model organisms and reuse the method for eventual genomic modifications makes this a valuable addition to the genetic engineering toolbox.

## Acknowledgments

L.T.R is partially supported by the Florida education fund McKnight doctoral fellowship. We thank L. Schuster and C. Mosby-Haundrup for their helpful conversations and comments on the manuscript. We thank C. Mejia and C. Espinoza for their technical assistance.

## Author contributions

C.R.R. conceived the study and performed the initial plasmid cloning. L.T.R. performed the transposition experiments, phenotype and genotype assays, and data analysis. A.J.E. prepared the next-generation sequencing library, and L.T.R. graphed and analyzed the data. C.R.R. and L.T.R. drafted the manuscript. All authors approve of the manuscript.

## Materials and Methods

### Bacterial strains, plasmid maintenance, media, and growth conditions

The complete lists of strains, plasmids, and primers are provided in Tables S1-4. *E. coli* was grown in Luria broth Miller (LB-Miller, 10 g/L tryptone, 10 g/L NaCl, 5 g/L yeast extract) at 37 °C with the following antibiotic concentrations when required: 50 μg/ml kanamycin (Km), 20 μg/ml gentamycin (Gm), 100 μg/ml carbenicillin (Carb), 20 μg/ml tetracycline (Tet) or 50 μg/ ml hygromycin B (Hyg). *B. thailandensis* E264 (Bt), *P. putida* KT2440 (Pp), and *A. fabrum* C58 (Af) were grown in Luria broth-Lennox (LB-Lennox, 10 g/L tryptone, 5 g/L NaCl, 5 g/L yeast extract) at 30 °C (Pp, Af, Bt) or 37 °C (Bt), with the following antibiotics when necessary: 200 μg/ml Km (Pp, Af) or 250 μg/ ml Km (Bt), 100 μg/ ml Gm (Pp, Af, Bt), 20 μg/ ml Tet (Pp, Af, Bt), 100 μg/ml Hyg (Af), and 2,000 μg/ml Zeocin (Zeo) (Bt, Pp). Gene expression regulated under a lactose inducible promoter uses 1 mM final concentration Isopropyl ß-D-1-thiogalactopyranoside (IPTG) inducer in the media. Auxotrophic mutants were grown in M9 minimal agar, final concentration recipe as follows: 1X M9 salts (6 g/L Na_2_HPO_4_, 3 g/L KH_2_PO_4_, 1 g/L NH_4_Cl, 0.5 g/L NaCl), 2 mM MgSO_4_•7H_2_O, 0.1 mM CaCl_2_, 0.2% (w/v) glucose, 1.5% (w/v) agar, supplemented with 0.8 mM L-tryptophan, 0.4 mM L-arginine, 0.4 mM L-serine and antibiotics as needed. Transformations were recovered in super optimal broth (SOB, 0.5% (w/v) yeast extract, 2% (w/v) tryptone, 10 mM NaCl, 2.5 mM KCl, 20 mM MgSO_4_).

### Plasmid cloning

All primers are listed in Tables S3 and S4. Primer pairs for pCasTn and pDonor cloning PCRs are listed in Supplementary Table S6. Q5 2X MM (NEB) was used following the manufacturer’s protocol. All PCR reactions were DpnI digested for 30 min and purified with a PCR clean-up kit prior to assembly with NEBuilder-HiFi assembly mix, except for pRO16Sh5 which was blunt ligated using T7 ligase (NEB).The assemblies were transformed into chemically competent *E. coli* NEB 5-alpha (NEB) and plated onto medium with the cognate antibiotic. <u>pCasTn plasmid cloning:</u> ShCAST genes were PCR amplified from pHelper_ShCAST_sgRNA (Addgene #127921). The vector backbones were amplified from plasmids carrying an RK2 origin of replication with a Kanamycin resistance gene for pKSh2 (Addgene #167515) or Gentamicin for pKSh5 (Addgene #149463), and a pBBR1 oriV for the pBLlSh2 backbone (Addgene #149470). The pRO1600 Ts origin was amplified from pFlpe2^50^. <u>pDonor plasmid construction:</u> The sgRNA was amplified from pHelper_ShCAST_sgRNA (Addgene #127921), and the transposon was amplified from pDonor_ShCAST_kanR (Addgene #127924). The two linear fragments were phosphorylated with T4 Polynucleotide Kinase and ligated with T4 DNA ligase, following the manufacturer’s protocols. <u>Violacein pDonor (pUCSh5vio-2300) cloning:</u> The plasmid backbone was amplified from pUCSh5-2300 (this study), and the violacein insert, the P_Llac_ promoter and *lacI* repressor were amplified from pBLlVio^52^. <u>pDonor re-targeting:</u> The pDonor sgRNA was re-targeted using around-the-horn cloning by amplifying the template with mRFP in place of the target sequence. The phosphorylated primer 2505 was paired with primers that possess overhangs containing the target sequence (Supplementary Table S4). The reaction was DpnI digested, purified with a silica column kit, ligated using T4 DNA ligase at room temperature (RT) overnight, and transformed into chemically competent *E. coli* cells. Colonies were screened the following day for the absence of red color, indicating loss of mRFP, and sequence verified.

### Creation of reporter strains with a yellow fluorescent protein

The *efyp* reporter strains of Bt, Pp, and Af were constructed with the mini-Tn7 system according to Choi et al^50^. The *efyp* integration in Bt was confirmed to be in chromosome 1 by PCR as described by Choi et al.^3^ (Supplementary Figure S2).

### Flp mediated marker excision in Bt and Pp

Bt and Pp were made electroporated with pFlpe2 Zeo (Pp or Bt) or pFlpe4 Km (Bt)^3^. After electroporation, Bt and Pp were recovered in SOB with 0.2% rhamnose to express *Flp* at 30 °C for 3 hrs, and the recovery was streaked out in an LB-Lennox agar 0.2% rhamnose with Zeo (Pp or Bt) or Km (Bt) and incubated at 30 °C. Colonies are then patched onto LB-Lennox plates with and without transposon antibiotic and incubated overnight, colonies that are antibiotic sensitive are grown in LB-Lennox for 2 hrs at 42 °C (Bt) or 37 °C (Pp) to cure the pFlpe2/pFlpe4 plasmids.

### CasTn Experiments

*<u>B. thailandensis</u>.* One small colony of Bt with pKSh5 or pRO16Sh5 was grown overnight at 30 °C 200 rpm in 5 mL LB Gm. The following day 4.5 mL of culture was spun down at 6,000 rpm for 3 min. The pellet was washed 2X with 300 mM sucrose at RT and resuspended in 300 μL of the sucrose solution. 50 μL of Bt resuspension was electroporated in a 0.1 cm cuvette at RT with 0.5 μL of pDonor plasmid (average ~200 ng) at 2.2 kV using a MicroPulser (Bio-Rad). The cells were recovered in 1 mL of SOB at 37 °C for 2 hrs, and 1:10 dilutions were plated on LB Km and incubated at 37 °C for 48 hrs until colonies appeared. *<u>P. putida</u>.* One small colony of Pp with pBLlSh2 was grown overnight at 30 °C 200 rpm in 5 mL LB Km, and the following day diluted to OD_600_ 0.1 in fresh LB Km and grown at 30 °C, 200 rpm. Upon reaching mid-log phase (OD_600_ 0.4-0.5), the culture was induced with 1 mM IPTG and grown for an additional 1 hr. The cells were then made electrocompetent with a 300 mM sucrose RT 2X wash and electroporated in a 0.1 cm cuvette at 1.8 kV, recovered in 1 mL SOB at 30 °C for 2 hrs and plated on LB Tet or Gm and incubated overnight at 30 °C until colonies appeared. For pRO16Sh2, a culture was struck out in LB-agar Km and grown overnight at 30 °C, the next day three colonies were passed into 5 mL LB Km 30 °C 200 rpm and grown until mid-log phase. At this time the cells were made electrocompetent and the protocol follows as previously described. *<u>A. fabrum</u>.* One colony of Af with pKSh2 was grown overnight in LB Km at 30 °C, 200rpm. The overnight culture was set back to an OD_600_ of 0.1 in 5 mL LB Km and then set back again at the mid-exponential phase. One mL of cells was then washed twice in 300 mM sucrose and then electroporated with pDonor at RT in a 0.1 cm cuvette with a 2.2 kV pulse. The cells were recovered in 1 mL SOB at 30 °C for 4 hrs, and 1:10 dilutions were plated on LB Hyg and incubated at 30 °C for 48 hrs. For all three bacteria, CFU counts were from an average of 32 colonies from each of three biological replicates that were picked into media plus antibiotic for selection of the transposon insertion mutants.

### Phenotypic Screens

Colonies obtained after transformation with pDonor were resuspended in sterile water and immediately spotted onto LB-agar with the appropriate antibiotic. The spots were resuspended in sterile water for loss of fluorescence screens, re-spotted onto an LB-agar plate, and incubated overnight at 30 °C. Plates were removed from the incubator and left at RT for one week to allow continued expression of the *eyfp* for better visualization. The plates were viewed through a blue light transilluminator (470 nm) and assessed for fluorescence. Loss of fluorescence on-target % was calculated as total_non-fluorescent mutants_/total_mutants_. For auxotrophy screens, each spot was resuspended in sterile water, and 5-10 μL was immediately spotted onto M9 agar with appropriate antibiotics with and without amino acid supplementation. The auxotrophy phenotype on-target % was calculated as total_auxotrophic_/ total_mutants_, total_auxotrophic_= #Mutants_M9+AA_ - #Mutants_M9_.

### Genotyping transposon insertion mutants

Supplementary Table S7 has primer pairs for the genotyping PCR reactions. An average of eight colonies per biological replicate and twenty-four total colonies per target were screened by colony PCR. KAPA2G Robust PCR Kit was used to make the PCR master mix using Buffer B with enhancer. Individual mutants were passed to a 96-well plate with 50 μL of water, and 1 μL of the cell suspension was passed to a PCR tube with the PCR reaction mix. Reactions were separated on a 2% agarose gel and imaged.

### Curing CasTn plasmids

CasTn plasmids were cured through overnight LB passages or growth at high temperature. Briefly one colony was passed into 1 ml of LB then 1 μl of the suspension was passed into 5 ml LB and grown overnight with shaking, the next day a second passage is done for Af. The next morning streaks are done in LB agar without antibiotics and grown overnight. Individial colonies are patched into LB with and without the antibiotic for the CasTn plasmid and incubated. For the Ts pCasTn the initial cell suspension is grown for 4 hrs at a higher temperature, 42 °C (Bt) or 37 °C (Pp), and then streaked out for patching following the same protocol as above. Antibiotc sensitive patches have lost the CasTn plasmid.

### TnSeq library preparation

Transposon insertion sites were mapped with TnSeq in independent integration experiments where the pDonor plasmid was transformed as described above, and the entire transformation was plated after two hours of recovery. The following day, all colonies in the plate were scraped into a 1.5 mL tube, washed, and resuspended with 250 μL of phosphate-buffered saline. Genomic DNA was then extracted using phenol:chloroform:isoamyl alcohol followed by ethanol precipitation^62^. Extracted gDNA was prepared for sequencing using a modified protocol^63^ for the NEBNext Ultra II FS DNA Library Prep Kit for Illumina (NEB) at ½ of the recommended reaction volume. After adaptor ligation, transposon-containing fragments were enriched by nested PCR amplification using primers 2625 and 2731 for 15 cycles, followed by primers 2761 and 2731 for 7 cycles. Samples were then PCR amplified for 7 cycles with Illumina adapters that possessed unique dual indexes using the NGS UDI Primer Set 48 (Eurofins Genomics). Sequencing was performed on an Illumina NovaSeq6000 (2 × 150 bp paired end reads, Novogene Co. Sacramento, CA). Bioinformatic analysis of on/off-targeting was performed by aligning trimmed and quality filtered reads to reference genomes using the Burrows-Wheeler Aligner v0.7.17^64^.

### TnSeq bioinformatic pipeline

Fastq reads were trimmed for the transposon right end and filtered for quality using the program cutadapt v3.2 (parameters: ctgatgacaataatttgtcacaacgacatata attagtcactgtaca -- discard-untrimmed -e 0.2 -q 16 --minimum-length=20). Reads that mapped to the backbone of the pDonor plasmid were then removed (parameters: gtagagacgtagcaatgctacctct ctacaatggttttgtatggtgcactctcag -- discard-trimmed -e 0.2). Subsequent cleaned reads were aligned to their respective bacteria reference file (.fn) using the Burrows-Wheeler Aligner (bwa) v0.7.17 with default parameters. GenBank accession or RefSeq numbers for reference genomes: *A. fabrum* (AE007869.2, AE007870.2, AE007871.2, AE007872.2), *B. thailandensis* (NC_007651.1, NC_007650.1) and *P. putida* (NC_002947.4). The Tn7 *eyfp* cassette was inserted in the genome sequence of each bacteria, with the RE towards *glmS*. It was inserted at nucleotide 1770721 of the circular chromosome of *A. fabrum*, nucleotide 6170446 of the *P. putida* chromosome, and at nucleotide 2360784 of *B. thailandensis* chromosome I. The bwa alignment file (.sam) output was checked for quality, sorted, and converted to a binary alignment file (.bam) using samtools v1.11. DeepTools v3.5 was used to create a .bedgraph file from the .bam file to map transposon insertion locations (bamCoverage parameters: -of bedgraph --normalizeUsing CPM --binSize 1000). Data were graphed using R^65^ v3.6.1, R studio v1.2.1335, and ggplot2^66^ package.

### Statistics

The efficiency, mean, and standard deviations were calculated using R^65^ v3.6.1, RStudio v1.2.1335, and table 1^67^ package. Comparisons of phenotype and genotype efficiency were made using Pearson correlation. Global statistical analysis between sgRNA length and efficiency was performed using the Kruskal-Wallis test. Statistical analysis of efficiency between paired groups of targets with different sgRNA lengths in Bt and Pp was done using the Mann-Whitney U test. At least three independent experiments were used for all tests, a p-value < 0.05 was considered significant.

### Data availability

Raw data for each mutant is in the supplementary dataset spreadsheet. Plasmid sequences, transformation efficiencies, transposon sequencing statistics, location of genomic targets, insertion distances, and main off-target regions are also in the supplementary dataset. Transposon sequencing data is available from the National Center for Biotechnology Information (NCBI) sequence read archive (SRA; BioProject accession: PRJNA756570). R script for data analysis can be found at Github (https://github.com/lidi-marie/ReischLab-CasTn).

