## Supplementary Information for "Broad-host-range mutagenesis with CRISPR-associated transposase"

**This PDF file includes:**

SI Text

Tables S1 to S14

Figures S1 to S17

Legends Datasets

SI References

**SI Text.**

**sgRNA target selection.** Targets were selected a Protospacer Adjacent Motif (PAM) of KGTB (K= G or T, B= T or C or G), within the first 300 bp from gene start. For auxotrophic targets, genes selected as targets for *P. putida* were based on published work by Molina-Henares et al. [1] on auxotrophic genes in M9 media. *B. thailandensis* and *A. fabrum* auxotrophic targets were selected judiciously to reduce possible polar effects from gene disruption.

**Supplementary Table S1 .** Strains used in this study**.**

| **Strain** | **Genotype and features** | **Reference** |
|---|---|---|
| *Agrobacterium fabrum* C58 | | |
| *Agrobacterium fabrum* C58 *eyfp* | $Gm^R$, *glmS* attTn7::miniTn7-*accC*1-*eyfp* | This work |
| *Burkholderia thailandensis* E264 | **Δ**(amrAB-oprA)::FRT | [2] |
| *Burkholderia thailandensis* E264 *eyfp* | **Δ**(amrAB-oprA)::FRT *glmS*1 attTn7::miniTn7-*eyfp* | This work |
| *Pseudomonas putida* KT2440 | | |
| *Pseudomonas putida* KT2440 *eyfp* | *glmS* attTn7::miniTn7- *eyfp* | This work |
| *Escherichia coli* NEB® 5-alpha | *fhuA2* (*argF-lacZ*)*U169 phoA glnV44 80* *(lacZ)*M15 *gyrA96 recA1 relA1 endA1 thi-1 hsdR17* | |

**Supplementary Table S2.** Plasmids used in this study.

| Plasmid | Features | Reference | Addgene no. |
|---|---|---|---|
| pKSh2 | $Km^R$ , RK2 ori, $P_{lac}$,Sh-*cas12K-tnsBC-tniQ-neo* | This work | 170979 |
| pKSh5 | $Gm^R$, RK2 ori, $P_{lac}$ ,Sh-*cas12K-tnsBC-tniQ -aaC1* | This work | 170980 |
| pBLlSh2 | $Km^R$, pBR322 ori, $P_{Llac}$, Sh-*cas12K-tnsBC-tniQ-neo* | This work | 170981 |
| pRO16Sh5 | $Gm^R$, pRO1600 Ts ori, $P_{lac}$ ,Sh-*cas12K-tnsBC-tniQ -aaC1* | This work | 180076 |
| pRO16Sh2 | $Km^R$ , pRO1600 Ts ori, $P_{lac}$,Sh-*cas12K-tnsBC-tniQ-neo* | This work | 180604 |
| pUC18T-mini-Tn7T-Gm-eyfp | $Amp^R$, $Gm^R$, ColE1 ori, Tn-*aacC1-eyfp* | [3] | |
| pUCSh2-mRFP | $Amp^R$, $Km^R$, ColE1 ori, *bla*, $P_{J23108}$-gRNA-*mRFP*, Tn-*neo* | This work | 170982 |
| pUCSh3-mRFP | $Amp^R$, $Tet^R$, ColE1 ori, *bla*, $P_{J23108}$-gRNA-*mRFP*, Tn-*tetA* | This work | 170983 |
| pUCSh8-mRFP | $Amp^R$, $Hyg^R$, ColE1 ori, *bla*, $P_{J23108}$-gRNA-*mRFP*, Tn-*hph* | This work | 171210 |
| pUCSh5vio-2300 | $Amp^R$, $Gm^R$, ColE1 ori, *bla*, $P_{J23108}$-sgRNA, Tn-*vioABCD-aacC1* | This work | 170987 |
| pFlpe2 | $Zeo^R$, ori1600-rep,oriT-ori, rhaS-rhaR-PrhaBAD-FLPe | [4] | |
| pFlpe4 | $Km^R$, $P_{S12}$-nptI-ori1600-rep, oriT-ori, rhaS-rhaR-PrhaBAD-FLPe | [4] | |
| pUCSh2-2300 | $Amp^R$, $Km^R$, ColE1 ori, *bla*, $P_{J23108}$-sgRNA, Tn-*neo* | This work | 170984 |
| pUCSh5-2300 | $Amp^R$, $Gm^R$, ColE1 ori, *bla*, $P_{J23108}$-sgRNA, Tn-*aacC1* | This work | 171788 |
| pUCSh3-2300 | $Amp^R$, $Tet^R$, ColE1 ori, *bla*, $P_{J23108}$-sgRNA, Tn-*tetA* | This work | 170985 |
| pUCSh8-2300 | $Amp^R$, $Hyg^R$, ColE1 ori, *bla*, $P_{J23108}$-sgRNA, Tn-*hph* | This work | 170986 |
| pUC18 | $Amp^R$ , ColE1 ori, *bla*, | [5] | |
| pHelper_ShCAST_sgRNA | $Amp^R$, ColE1 ori, Sh-*cas12K-tnsBC-tniQ*, $P_{J23119}$-gRNA | [6] | |
| pDonor_ShCAST_kanR | $Km^R$, R6K ori, ColE1 ori, Tn-*neo* | [6] | |

**Supplementary Table S3.** Oligoes used in this study.

| Oligo | Sequence (5’-3’) | Description/Reference |
|---|---|---|
| 2599BtArgCF | GCCGCGCGATCCGGCAAG | Gene primer |
| 2615AtCysF | CGAATGCGGTCTTCACGTCG | Gene primer |
| 2616AtLeuBF | GGACGGAACGAGTTTGACG | Gene primer |
| 2548AtcysDR | GGAATGTTTTCCGCCTCGATGT | Gene primer |
| 2555BttrpAR | ATAGTAGACGTAACCGCTCGCC | Gene primer |
| 2562PpserAR | GAACTCTTCGTCGTTGGAGCGT | Gene primer |
| 2618BtTrpAR2 | GCGAAGCGTCACGCCTTTC | Gene primer |
| 2619ShLE | GGTGGGTTGAAAGCAAGTCC | LE primer upstream |
| 2910motAF | CAACCCGTCGAGATCCTGATGA | Gene primer |
| 2919B-lactR | GCAAGCCCGTCAATCCGATG | Gene primer |
| 2626ShRE | CCTGCCACTCATCGCAGTCTAGCTTGGATT | RE primer upstream |
| 2624ShLEseq | CAACGCTGATGGGTCACGAC | LE sequencing primer |
| 2625ShREseq | GTCACTGCAAAGAGGTTATGC | RE sequencing primer |
| 2630PpTrpAF | GTTGCCCAAGCGTTACATCG | Gene Primer |
| 2631PpSerAF | AGCACTCTTCACCTGTCGGG | Gene primer |
| 75 | ATTGCTACAGGCATCGTGGT | pUC backbone PCR |
| 1531 | AATGTGCGCGGAACCCC | pUC backbone PCR |
| 1535 | GACGAAAGGGCCTCGTGA | sgRNA sequencing |
| 1482glmS1-DN5 | GTTCGTCGTCCACTGGGATCA | Bt *glmS*-1 gene primer[4] |
| 1483glmS2-DN5 | AGATCGGATGGAATTCGTGGAG | Bt *glmS*-2 gene primer[4] |
| 1484primerTn7L | ATTAGCTTACGACGCTACACCC | Mini-Tn7 LE primer[4] |
| 1857 | GTCCAGATAGCCCAGTAGC | LE primer |
| 1370 | ATATCCAATTGCCTGCCACTCATCGCAGTCTAGCTTGGATT | RE primer |
| 2731PalanitoU7 | GTTCAGACGTGTGCTCTTCCGATCTGTCTCGTGGGCTCGGA GATGTGTATAAG | TnSeq |
| 2761REadapter | CCTACACGACGCTCTTCCGATCTCTGATGACAATAATTTGTC ACAACGAC | TnSeq |
| 1337 | GTACAGATATCACTAGTGAGCTCATGCATG | WPI* |
| 2216 | GCCTGCAGTCTAGACTCGAGCTGATACCGCTCGCCGCAG | WPI* |
| 1537 | TAAGCTTAATTAGCTGAGCTTGG | WPI* |
| 2735 | AGCGAAACGATTTGGAACCAGT | WPI* |
| 2220 | GGTTTCCCGACTGGAAAGCGGGGTTCCGCGCACATTTC | pDonor cloning |
| 1969 | GTAAGGATCTCCAGGCATC | pDonor cloning |
| 2368 | ATCACGAGGCCCTTTCGTCAATGACCCCGAAGCAGGG | pDonor cloning |
| 2503 | TCCCCGAAAAGTGCCACCCAGGATTAGCAGAGCGAG | pDonor cloning |
| 1494 | TGCGCTCTTCCCTGTCCGCTTCCTCGCTCACT | pDonor cloning |
| 1530 | CAGGTGGCACTTTTCGGGGAAATGTGCGCGGAACCCC | pDonor cloning |
| 2221 | GTAAAACGACGGCCAGTGAGCTGGCCTTTTGCTCAACG | pCasTn cloning |
| 2393 | GTTTCCCGACTGGAAAGCGTAGAGAGCGTTCACCG | pCasTn cloning |
| 1108 | TCACTGGCCGTCGTTTTACAACGTCGTGAC | pCasTn cloning |
| 2822 | GCTTTCCAGTCGGGAAACCTG | pCasTn cloning |
| 2854 | TCACTGGCCGTCGTTTTACAACAGCTCTGCAGATTTCGT | pRO16Sh cloning |
| 2855 | GGTGGCACTTTTCGGGGAACGCAGGAAAGAACATGGGG | pRO16Sh cloning |
| 1529 | TCCCCGAAAAGTGCCACC | pRO16Sh cloning |

*** WPI = Whole plasmid integration**

**Supplementary Table S4.** pDonor sgRNA target cloning oligoes.

| Oligo | Sequence (5'-3') |
|---|---|
| 2505ShRPO4 | CTTTCAACCCATTTAGGGTTCC |
| 2508At_cysD_tgtc | GCATCCAAGCCGCCGCTCGATCCGgctcaccttcgggtgg |
| 2509At_cysD_ggtt | ATCGAATTCGGCGGCGACTTCGCGgctcaccttcgggtgg |
| 2511At_leuB_tgtt | CGGTGCCGTTGGCGGCCCGAAATGgctcaccttcgggtgg |
| 2516Bt_argC_TGTC | GACGGCCAGGAAGGCACGACCGGTgctcaccttcgggtgg |
| 2517Bt_argC_GGTC | TGAAGATCTTCGAATATCTGTCCGgctcaccttcgggtgg |
| 2523Bt_trpA_GGTC | GAGCTGATGCACGCGCTTGCCGAAgctcaccttcgggtgg |
| 2524Bt_trpA_GGTC | CCCCGCGGTGATGAACGGGATCAGgctcaccttcgggtgg |
| 2541Pp_serA_GGTT | TACGCAGATGAGCAAGACTTCTCTgctcaccttcgggtgg |
| 2542Pp_serA_TGTT | GGTGTAGCCGGCGGCCTTGAGGGTgctcaccttcgggtgg |
| 2544Pp_trpF_GGTT | CTTGTTCCATGAGCAATGTTCGCAgctcaccttcgggtgg |
| 2545Pp_trpF_TGTT | CGCAGCAAGATCTGCGGGATTACCgctcaccttcgggtgg |
| 2621BtTrpA3cgtg | CCGTTCTCCGATCCGATGGCCGACgctcaccttcgggtgg |
| 2622BtTrpA4ggtc | GCGCTCGCGAAAGCGCCTCACGTCgctcaccttcgggtgg |
| 2655Pp2542_20bp | GGTGTAGCCGGCGGCCTTGAgctcaccttcgggtgg |
| 2656Pp2542_28bp | GGTGTAGCCGGCGGCCTTGAGGGTATCGgctcaccttcgggtgg |
| 2657Bt2517_20bp | TGAAGATCTTCGAATATCTGgctcaccttcgggtgg |
| 2658Bt2517_28bp | TGAAGATCTTCGAATATCTGTCCGCGCGgctcaccttcgggtgg |
| 2664Bt2523_16bp | GAGCTGATGCACGCGCgctcaccttcgggtgg |
| 2300eypfGGTC2F | GGGTAGCGGGCGAAGCACTGCAGgctcaccttcgggtgg |
| 2686eyfpGGTC17 | GAGCTGGACGGCGACGTgctcaccttcgggtgg |
| 26872300GGTC17bp | GGGGTAGCGGGCGAAGCgctcaccttcgggtgg |
| 2688eyfpGGTT17bp | CACCAGGGTGTCGCCCTgctcaccttcgggtgg |
| 27082686_24bp | GAGCTGGACGGCGACGTAAACGGCgctcaccttcgggtgg |
| 2714AtleuBtgtt17 | CGGTGCCGTTGGCGGCCgctcaccttcgggtgg |
| 2885Motility | GTCGATGTCCGCCTCGAGCGTGAGgctcaccttcgggtgg |
| 2887Beta-lact | GACTCGAGTTCGCGCAATTGCTGCgctcaccttcgggtgg |

**Supplementary Table S5.** sgRNA target sequence and target identifier used in this study.

| Target name | Cloning oligo name | Target Sequence (5'-3') |
|---|---|---|
| ***cysD*-1** | 2508At_cysD_tgtc24bp | GCATCCAAGCCGCCGCTCGATCCG |
| ***cysD*-2** | 2509At_cysD_ggtt24bp | ATCGAATTCGGCGGCGACTTCGCG |
| ***leuB*-1** | 2511At_leuB_tgtt24bp | CGGTGCCGTTGGCGGCCCGAAATG |
| ***argC*-1** | 2516Bt_argC_TGTC24bp | GACGGCCAGGAAGGCACGACCGGT |
| ***argC*-2** | 2517Bt_argC_GGTC24bp | TGAAGATCTTCGAATATCTGTCCG |
| ***trpA*-1** | 2523Bt_trpA_GGTC24bp | GAGCTGATGCACGCGCTTGCCGAA |
| ***trpA*-2** | 2524Bt_trpA_GGTC24bp | CCCCGCGGTGATGAACGGGATCAG |
| ***serA*-1** | 2541Pp_serA_GGTT24bp | TACGCAGATGAGCAAGACTTCTCT |
| ***serA*-2** | 2542Pp_serA_TGTT24bp | GGTGTAGCCGGCGGCCTTGAGGGT |
| ***serA-3*** | 2543Pp_serA_GGTC24bp | AACCTGGTTGGTGCCGATGCAGAA |
| ***trpF*-1** | 2544Pp_trpF_GGTT24bp | CTTGTTCCATGAGCAATGTTCGCA |
| ***trpF*-2** | 2545Pp_trpF_TGTT 24bp | CGCAGCAAGATCTGCGGGATTACC |
| ***trpA*-3** | 2621BtTrpA3cgtg24bp | CCGTTCTCCGATCCGATGGCCGAC |
| ***trpA*-4** | 2622BtTrpA4ggtc24bp | GCGCTCGCGAAAGCGCCTCACGTC |
| ***serA*-4** | 2655Pp2542_20bp | GGTGTAGCCGGCGGCCTTGA |
| ***serA*-5** | 2656Pp2542_28bp | GGTGTAGCCGGCGGCCTTGAGGGTATCG |
| ***argC*-3** | 2657Bt2517_20bp | TGAAGATCTTCGAATATCTG |
| ***argC*-4** | 2658Bt2517_28bp | TGAAGATCTTCGAATATCTGTCCGCGCG |
| ***trpA*-5** | 2664Bt2523_16bp | GAGCTGATGCACGCGC |
| ***eyfp*-1** | 2300eypfGGTC24bp | GGGGTAGCGGGCGAAGCACTGCAG |
| ***eyfp*-2** | 2686eyfpGGTC17bp | GAGCTGGACGGCGACGT |
| ***eyfp*-3** | 26872300GGTC17bp | GGGGTAGCGGGCGAAGC |
| ***eyfp*-4** | 2688eyfpGGTT17bp | CACCAGGGTGTCGCCCT |
| ***eyfp*-5** | 27082686_24bp | GAGCTGGACGGCGACGTAAACGGC |
| ***leuB*-2** | 2714AtleuBtgtt17 | CGGTGCCGTTGGCGGCC |
| ***motA-1*** | 2885Motility | TCGATGTCCGCCTCGAGCGTGAG |
| **β-lac-1** | 2887Beta-lact | GACTCGAGTTCGCGCAATTGCTGC |

**Supplementary Table S6.** PCR reactions used for cloning pCasTns and pDonors.

| Purpose | Template | Primers | |
|---|---|---|---|
| pCasTn cloning - ShCAST genes | pHelper_ShCAST_sgRNA | 2221 | 2393 |
| pCasTn cloning- RK2 ori | pK plasmids | 1108 | 2822 |
| pDonor cloning- sgRNA | pHelper_ShCAST_sgRNA | 2220 | 1969 |
| pDonor cloning- transposon | pDonor_ShCAST_kanR | 2368 | 2503 |
| pDonor cloning- pUC backbone | pUC18 | 1494 | 1530 |
| pRO16Sh5 cloning - pRO1600 Ts origin | pFlpe2[4] | 2855 | 2856 |
| pRO16Sh5 cloning- ShCAST genes | pKSh5 | 1529 | 2221 |
| pRO16Sh2 cloning - Origing + ShCAST | pRO16Sh5 | 2393 | 2854 |
| pRO16Sh2 cloning - Kan marker | pKSh2 | 1529 | 2822 |

**Supplementary Table S7.** PCR reactions for whole plasmid integrations and transposon insertion genotyping.

| PCR | Bacteria | Purpose | Target | Primers | | Expected Size (bp)‡ | Comment |
|---|---|---|---|---|---|---|---|
| **Backbone** | Af, Pp, Bt | WPI* | pDonor ori | 1531 | 75 | 662 | *Whole plasmid integration (WPI) |
| **RE-eyfp-1** | Af, Pp, Bt | RE | *eyfp-1* | 2626 | 2220 | 1092/747(off/on-target) | |
| **LE-eyfp-2** | Af, Pp, Bt | LE | *eyfp-2* | 2619 | 2220 | 548 | |
| **RE-eyfp-3** | Af, Pp, Bt | RE | *eyfp-3* | 1370 | 2220 | 1092/747(off/on-target) | |
| **RE-eyfp-4** | Af, Pp, Bt | RE | *eyfp-4* | 1370 | 2220 | 1092/882(off/on-target) | |
| **LE-eyfp-5** | Af, Pp, Bt | LE | *eyfp-5* | 2619 | 2220 | 548 | |
| **RE-serA-1** | Pp | RE | *serA-1* | 1370 | 2631 | 544 | |
| **LE-serA-2** | Pp | LE | *serA-2* | 1857 | 2562 | 1261 | |
| **LE-serA-3** | Pp | LE | *serA-3* | 1857 | 2562 | 1,092 | |
| **LE-serA-4** | Pp | LE | *serA-4* | 1857 | 2562 | 1261 | short *serA*-2 |
| **LE-serA-5** | Pp | LE | *serA-5* | 1857 | 2562 | 1261 | Long *serA*-2 |
| **LE-trpF-1** | Pp | LE | *trpF-1* | 1857 | 2630 | 641 | |
| **LE-trpF-2** | Pp | LE | *trpF-2* | 1857 | 2630 | 663 | |
| **LE-argC-1** | Bt | LE | *argC-1* | 2599 | 2619 | 418 | |
| **LE-argC-2** | Bt | LE | *argC-2* | 2599 | 2619 | 443 | |
| **LE-argC-3** | Bt | LE | *argC-3* | 2599 | 1857 | 721 | short *argC*-2 |
| **LE-argC-4** | Bt | LE | *argC-4* | 2599 | 1857 | 721 | long *argC*-2 |
| **RE-trpA-1** | Bt | RE | *trpA-1* | 1370 | 2555 | 717 | |
| **LE-trpA-2** | Bt | LE | *trpA-2* | 2619 | 2555 | 700 | |
| **LE-trpA-3** | Bt | LE | *trpA-3* | 2735 | 1857 | 707 | |
| **RE-trpA-4** | Bt | RE | *trpA-4* | 1370 | 2555 | 587 | |
| **RE-trpA-5** | Bt | RE | *trpA-5* | 1370 | 2555 | 717 | short *trpA*-1 |
| **RE-motA-1** | Bt | RE | *motA-1* | 2626 | 2910 | 510 | |
| **RE-β-lac-1** | Bt | RE | β-lac-1 | 2626 | 2919 | 700 | |
| **LE-cysD-1** | Af | LE | *cysD-1* | 2615 | 1857 | 665 | |
| **LE-cysD-2** | Af | LE | *cysD-2* | 2548 | 2619 | 777 | |
| **LE-leuB-1** | Af | LE | *leuB-1* | 2619 | 2616 | 548 | |
| **LE-leuB-2** | Af | LE | *leuB-2* | 2619 | 2616 | 548 | |
| **PCR-A** | Bt | WPI* | Flp mutant | 1337 | 2216 | 2191 | **Supplementary Figure S10.** |
| **PCR-B** | Bt | WPI* | Flp mutant | 2625 | 2619 | 2928 | **Supplementary Figure S10.** |
| **PCR-C** | Bt | WPI* | Flp mutant | 2300 | 1537 | 2221 | **Supplementary Figure S10.** |
| **PCR-D** | Bt | WPI* | Flp mutant | 2735 | 2618 | 260(WT),2470(NI), 1134(FO),7306(WPI)** | **WT= wild-type, NI= normal integration, FO= flipped out. **Fig S10.** |

‡Assuming 65 bp insertion distance from PAM for transposon insertion left-end or right-end (LE/RE) PCR.

**Supplementary Table S8.** Transformation efficiencies for Proteobacteria by CasTn system.

| | *A. fabrum* | | *B. thailandensis* | | *P. putida* | |
|---|---|---|---|---|---|---|
| | **pBLlSh2 (N=20)** | **pKSh2 (N=56)** | **pKSh5 (N=81)** | **pRO16Sh5 (N=9)** | **pBLlSh2 (N=32)** | **pRO16Sh2 (N=12)** |
| **Transformation efficiency (CFU/µg DNA)*** | | | | | | |
| **Mean (SD)** | 3.9E+04 (±4.6 E+04) | 2.0E+04 (±2.2E+04) | 8.3E+03 (±3.4E+04) | 4.8E+05 (±4.4E+05) | 2.5E+04 (±4.2E+04) | 1.2E+05 (±1.9E+05) |
| **Median [Min,Max]** | 1.5 E+04 [0,1.6E+05] | 1.3E+04 [0,8.0E+04] | 1.8E+03 [0,2.9E+05] | 1.4E+05 [7.1E+04, 1.2E+06] | 9.0E+03 [0,1.8E+05] | 2.8E+04 [62.9, 5.4E+05] |

*Transformation efficiency inclusive for all targets per CasTn system. Colony Forming Unit (CFU) per µg DNA transformed.

**Supplementary Table S9.** Global statistics genotype efficiency vs different length sgRNA.

| **Kruskal-Wallis rank sum test** | **H** | **df** | **p value‡** |
|---|---|---|---|
| *Agrobacterium fabrum* C58 *eyfp* | 2.90 | 1 | 0.088 |
| *Burkholderia thailandensis* E264 *eyfp* | 0.891 | 4 | 0.926 |
| *Pseudomonas putida* KT2440 *eyfp* | 0.980 | 2 | 0.613 |
| **Across all bacteria** | 1.71 | 4 | 0.789 |

‡ Significance codes: 0 ‘***’ 0.001 ‘**’ 0.01 ‘*’ 0.05 , alpha = 0.05
H= test statistic, df= degrees of freedom

**Supplementary Table S10.** Pearson correlation of genotype vs phenotype efficiency.

| Pearson correlation | t | df | CI 95% | Cor | p value‡ |
|---|---|---|---|---|---|
| *Agrobacterium fabrum* C58 *eyfp* | 9.898 | 5 | 0.8377,0.9965 | 0.9754 | 0.00018 *** |
| *Burkholderia thailandensis* E264 *eyfp* | 23.42 | 14 | 0.9633,0.9958 | 0.9875 | 1.25e-12 *** |
| *Pseudomonas putida* KT2440 *eyfp* | 15.31 | 13 | 0.9198, 0.9913 | 0.9734 | 1.06e-09 *** |
| **Across all bacteria** | 31.42 | 36 | 0.9658 , 0.9908 | 0.9823 | <2e-16 *** |

‡ Significance codes: 0 '***' 0.001 '**' 0.01 '*' 0.05 , alpha = 0.05
t = t-test statistic, df= degrees of freedom, CI 95%= 95% confidence interval,
Cor = Pearson correlation

**Supplementary Table S11.** Transformation efficiencies of *A. fabrum* by sgRNA target.

| sgRNA Target | Transformation efficiency (CFU/µg DNA)* | sgRNA Target | Transformation efficiency (CFU/µg DNA)* |
|---|---|---|---|
| ***eyfp*-1 (N=14)** | Mean (SD): 2.8E+04 (±3.0E+04)<br>Median [Min, Max]:<br>2.0E+04 [2.7E+03, 1.1E+05] | ***eyfp*-2 (N=6)** | Mean (SD): 4.3E+04 (±2.9E+04)<br>Median [Min, Max]:<br>5.3E+04 [5.0E+03, 7.1E+04] |
| ***cysD*-1 (N=3)** | Mean (SD): 1.1E+03 (±9.3E+02)<br>Median [Min, Max]:<br>1.4E+03 [0, 1.8E+03] | ***eyfp*-3 (N=6)** | Mean (SD): 4.0E+04 (±4.1E+04)<br>Median [Min, Max]:<br>3.1E+04 [7.1E+04, 1.1E+05] |
| ***cysD*-2 (N=6)** | Mean (SD): 1.4 E+04 (±1.6E+04)<br>Median [Min, Max]:<br>8.4E+03 [1.2E+02, 3.9E+04] | ***eyfp*-4 (N=6)** | Mean (SD): 4.9E+04 (±5.7E+04)<br>Median [Min, Max]:<br>2.9E+04 [1.1E+04, 1.6E+05] |
| ***leuB*-1 (N=5)** | Mean (SD): 9.5E+03 (±9.7E+03)<br>Median [Min, Max]:<br>5.0E+03 [1.4E+03, 2.3E+04] | ***eyfp-5* (N=3)** | Mean (SD): 4.9E+04 (±2.7E+04)<br>Median [Min, Max]:<br>3.5E+04 [3.1E+04, 8.0E+04] |

*Transformation efficiency by sgRNA target. Colony Forming Unit (CFU) per µg DNA transformed.

**Supplementary Table S12.** Transformation efficiencies of *B. thailandensis* by sgRNA target.

| sgRNA Target | Transformation efficiency (CFU/µg DNA)* |
|---|---|
| ***eyfp*-1 (N=3)** | Mean (SD): 1.5E+03 (±3.0E+02)<br>Median [Min, Max]:<br>1.4E+03 [1.2E+03, 1.8E+03] |
| ***argC*-1 (N=8)** | Mean (SD): 2.1E+03 (±1.3E+03)<br>Median [Min, Max]:<br>1.9E+03 [6.7E+02, 4.0 E+03] |
| ***argC*-2 (N=5)** | Mean (SD): 4.8E+03 (±2.9E+03)<br>Median [Min, Max]:<br>5.0E+03 [2.0E+03, 8.4E+03] |
| ***trpA*-1 (N=3)** | Mean (SD): 3.7E+04 (±5.1 E+04)<br>Median [Min, Max]:<br>7.4E+03 [7.2E+03, 9.6E+04] |
| ***trpA*-2 (N=6)** | Mean (SD): 2.0E+03 (±2.6E+03)<br>Median [Min, Max]:<br>4.9E+02 [0, 5.6E+03] |
| ***trpA*-3 (N=3)** | Mean (SD): 1.7E+03 (±1.2E+03)<br>Median [Min, Max]:<br>1.7E+03 [4.3E+02, 2.8E+03] |
| ***trpA*-4 (N=3)** | Mean (SD): 1.6E+03 (±1.2E+03)<br>Median [Min, Max]:<br>1.5E+03 [478, 2.8E+03] |
| ***argC*-3 (N=3)** | Mean (SD): 2.4E+03 (±1.7E+03)<br>Median [Min, Max]:<br>1.6E+03 [1.4E+03, 4.2E+03] |
| ***argC*-4 (N=2)** | Mean (SD): 2.8E+03 (±8.3E+02)<br>Median [Min, Max]:<br>2.8E+03 [2.3E+03, 3.4E+03] |
| ***trpA*-5 (N=3)** | Mean (SD): 3.8Ee+03 (±2.4E+03)<br>Median [Min, Max]:<br>3.0E+03 [1.8E+03, 6.5E+03] |
| ***eyfp*-2 (N=3)** | Mean (SD): 5.7E+03 (±4.1E+03)<br>Median [Min, Max]:<br>5.7E+03 [1.7E+03, 9.8E+03] |
| **Double Target *eyfp*-2 *trpA*-1 (N=3)** | Mean (SD): 1.2E+05 (±1.5E+05)<br>Median [Min, Max]:<br>5.0E+04 [9.7E+03, 2.9E+05] |
| ***eyfp*-3 (N=3)** | Mean (SD): 2.1E+03 (±1.9E+03)<br>Median [Min, Max]:<br>1.9E+03 [3.8E+02, 4.1E+03] |
| ***eyfp*-5 (N=3)** | Mean (SD): 7.7E+03 (±7.9E+03)<br>Median [Min, Max]:<br>6.1E+03 [6.7E+02, 1.6E+04] |

*Transformation efficiency (TE) by sgRNA target. Colony Forming Unit (CFU) per µg DNA transformed. TE of *eyfp*-4 not collected.

**Supplementary Table S13.** Transformation efficiencies of *P. putida* by sgRNA target.

| sgRNA Target | Transformation efficiency (CFU/µg DNA)* |
|---|---|
| ***eyfp*-1 (N=6)** | Mean (SD): 1.9E+04 (±3.0E+04)<br>Median [Min, Max]:<br>5.1E+03 [2.5E+03, 7.9E+04] |
| ***serA*-1 (N=3)** | Mean (SD): 6.8E+04 (±1.0E+05)<br>Median [Min, Max]:<br>1.4E+04 [7.6E+03, 1.8E+05] |
| ***serA-2* (N=4)** | Mean (SD): 2.8E+04 (±5.3E+04)<br>Median [Min, Max]:<br>2.7E+03 [0, 1.1E+05] |
| ***serA-3* (N=3)** | Mean (SD): 6.9E+03 (±9.9E+03)<br>Median [Min, Max]:<br>1.7E+03 [5.3E+02, 1.8E+04] |
| ***serA-4* (N=3)** | Mean (SD): 2.4E+03 (1.6E+03)<br>Median [Min, Max]:<br>3.1E+04 [579, 3.6E+03] |
| ***serA-5* (N=3)** | Mean (SD): 1.5E+03 (1.1E+03)<br>Median [Min, Max]:<br>2.1E+03 [247, 2.2E+03] |
| ***trpF*-1 (N=4)** | Mean (SD): 3.5E+04 (±3.9E+04)<br>Median [Min, Max]:<br>2.3E+04 [2.7E+03, 9.0E+03] |
| ***trpF*-2 (N=3)** | Mean (SD): 2.1E+04 (±6.9E+03)<br>Median [Min, Max]:<br>2.1E+04 [1.4E+04, 2.7E+04] |
| ***eyfp*-2 (N=4)** | Mean (SD): 2.1E+04 (3.8E+04)<br>Median [Min, Max]:<br>2.2E+03 [733, 7.7E+04] |
| ***eyfp-3* (N=3)** | Mean (SD): 1.2E+03 (212)<br>Median [Min, Max]:<br>1.1E+03 [1.1E+03, 1.5E+03] |
| ***eyfp-4* (N=3)** | Mean (SD): 2.4E+03 (1.9E+03)<br>Median [Min, Max]:<br>3.2E+04 [270, 3.8E+03] |
| ***eyfp*-5 (N=4)** | Mean (SD): 1.8E+04 (2.9E+04)<br>Median [Min, Max]:<br>3.4E+03 [3.1E+03, 6.1E+04] |
| ***eyfp*-1 Violacein (N=3)** | Mean (SD): 7.7E+03 (±1.3E+04)<br>Median [Min, Max]:<br>1.8E+02 [0, 2.3E+04] |
| ***eyfp-1* Gent (N=3)** | Mean (SD): 2.4E+04 (1.1E+04)<br>Median [Min, Max]:<br>2.4E+04 [1.4E+04, 3.6E+04] |

*Transformation efficiency by sgRNA target. Colony Forming Unit (CFU) per µg DNA transformed.

**Supplementary Table S14.** Genotyping results for cointegrate insertion (%).

| | ***Agrobacterium fabrum*** **(N=27)** | ***Burkholderia thailandensis*** **(N=52)** | ***Pseudomonas putida*** **(N=42)** | **Overall (N=121)** |
|---|---|---|---|---|
| Cointegrate insertion percent* | | | | |
| Mean (SD) | 80.5 (25.5) | 43.4 (29.5) | 31.6 (35.0) | 47.4 (35.7) |
| Median [Min, Max] | 96.9 [31.3, 100] | 42.9 [0, 100] | 13.4 [0, 100.0] | 42.9 [0, 100] |

*Cointegrate occurrence as mean of plasmid insertions per sgRNA target, biological replicates (N).

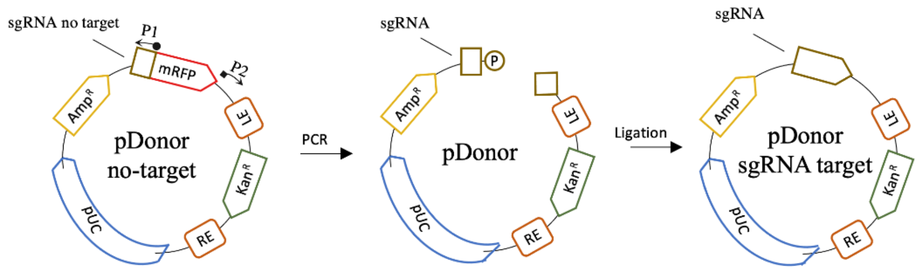


**Supplementary Figure S1. pDonor re-targeting through around-the-horn cloning.** Re-targeting was done through PCR with a 5'-phosphorylated reverse primer (P1) and a forward primer with a 5' overhang with the new target sequence (P2), leaving out mRFP from the amplicon. After PCR, the amplicon is ligated and transformed in *E.coli* for maintenance and retrieval. Red fluorescent *E. coli* colonies signify cloning failure.

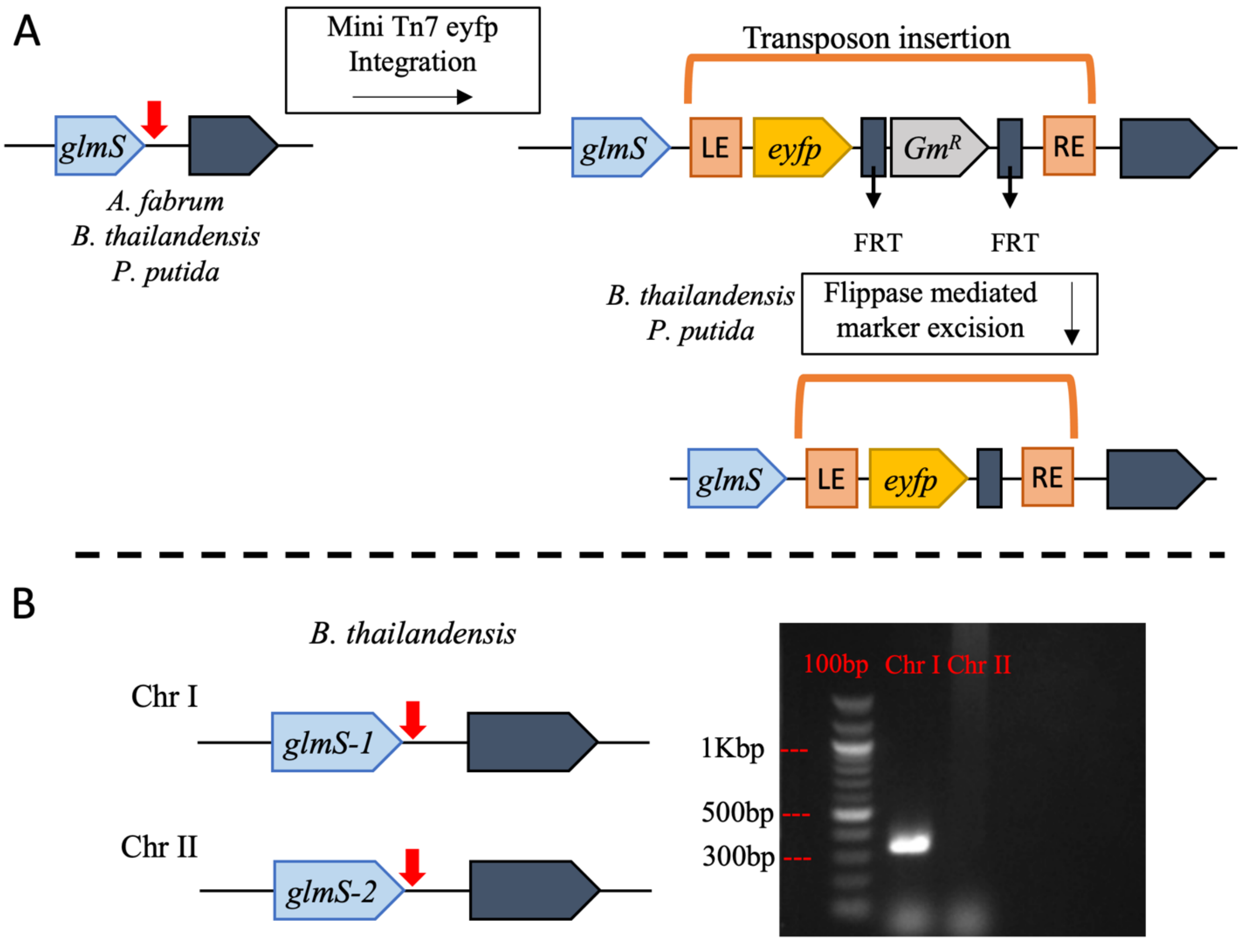


**Supplementary Figure S2. Insertion of *eyfp* cassette across targeted Proteobacteria through a mini Tn7 system. (A)** A cassette carrying *eyfp* and gentamycin (Gm) resistance was integrated in AT, BT, and PP, through a site-specific mini Tn7 system that recognizes a conserved attachment site (red arrow) found downstream of *glmS* (D-fructose-6-phosphate aminotransferase). The Gm marker is flanked by flippase recognition target (FRT) sites that allow excision using a flippase system (pFlpe2), in BT and PP. **(B)** Schematic of BT chromosomes carrying two copies of *glmS* both with attachment sites, *glmS*-1 and *glmS*-2 in chromosome 1 (Chr 1) and chromosome 2 (Chr 2) respectively. Colony PCR amplification was done with chromosome specific primers and a transposon LE primer with an expected band size at 322 bp (Chr 1) and 228 bp (Chr 2) ran on a 2% agarose gel with 100 bp DNA ladder.

A

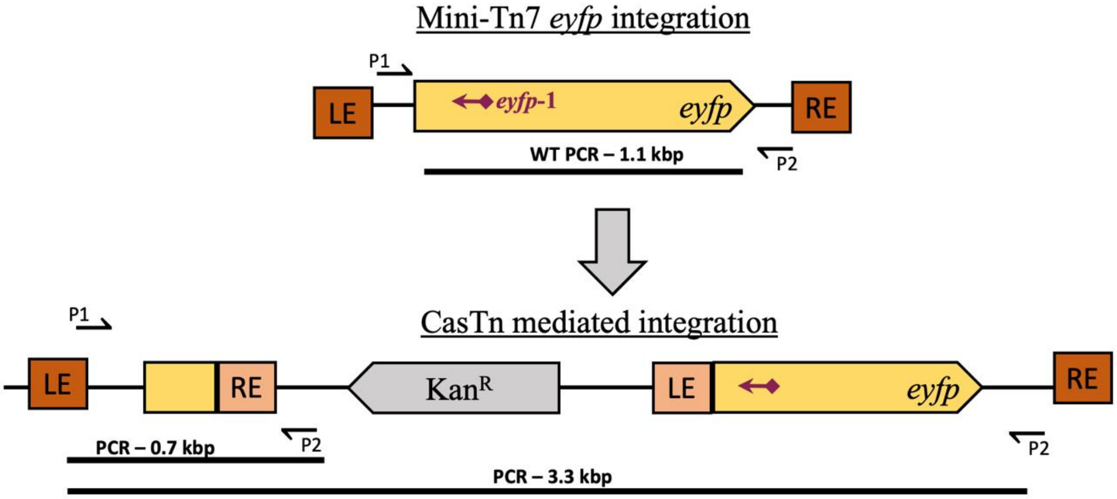


B

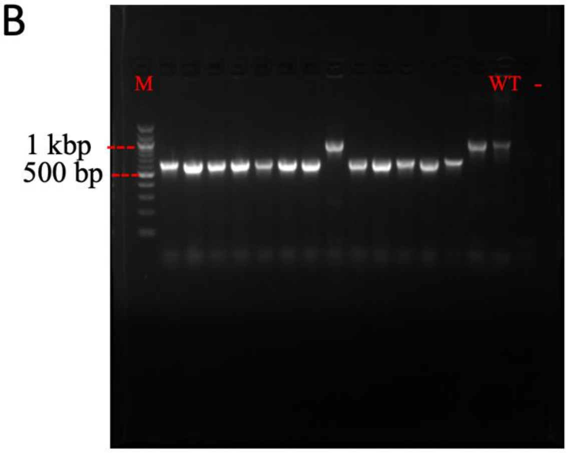


**Supplementary Figure S3. Yellow fluorescence CasTn targeting and insertion genotyping. (A)** CasTn targeted transposon integration using target *eyfp*-1. PCR of WT bacteria with the *eyfp* integration using primers P1 and P2 results in a 1.1 Kbp band. Successful integration into the target site allows PCR amplicon size change to ~700 bp. **(B)** PCR amplification of *eyfp*-1 targets in *B. thailandensis* transposon mutants separated on a 2% agarose gel with 100 bp DNA ladder (M), using *B. thailandensis* pKSh5 (WT) and a no template (-) controls.

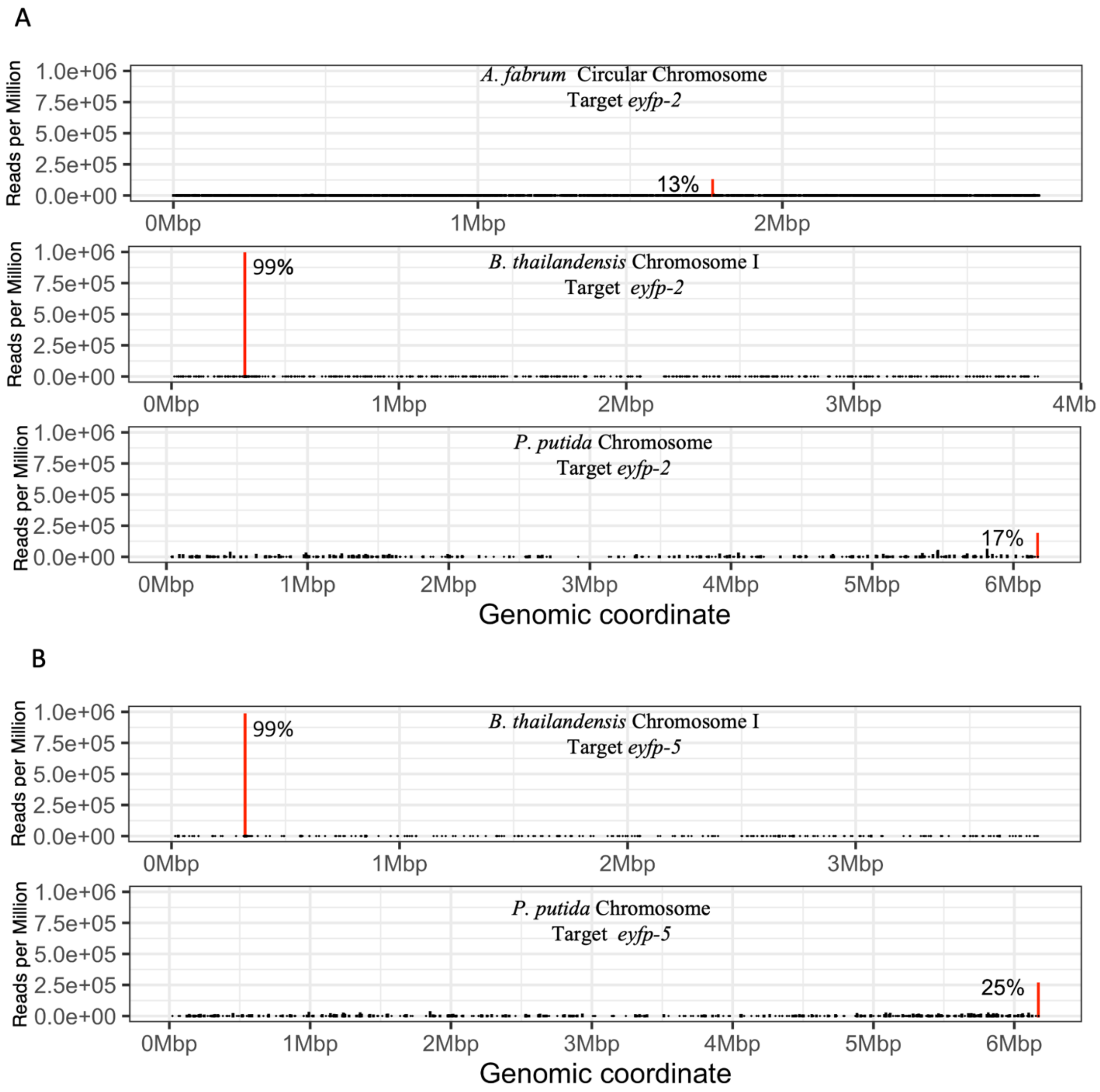


**Supplementary Figure S4. TnSeq data for *eyfp* targets *eyfp*-2 and *eyfp*-5 across Proteobacteria.** Transposon sequencing results for *eyfp* targets *eyfp*-2 **(A)** and *eyfp*-5 **(B)** in reads per million (RPM) for *A. fabrum*, *B. thailandensis*, and *P. putida***.** On-target peaks (red) are labeled with the percentage of reads mapped across the whole genome.

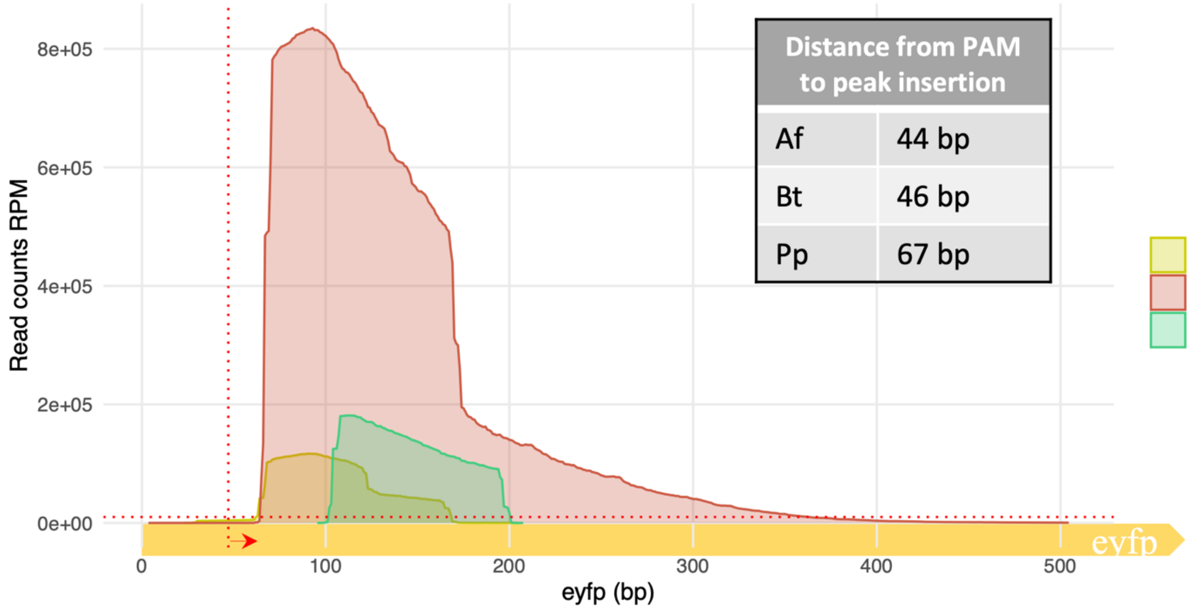


**Supplementary Figure S5. TnSeq comparison of target *eyfp*-2 transposon insertions.** Density plot of transposon insertions for target *eyfp*-2 in *Agrobacterium fabrum* (Af), *Burkholderia thailandensis* (Bt) and *Pseudomonas putida* (Pp) in reads per million (RPM). The x-axis goes from the start of *eyfp* (0 bp) to 500 bp into the gene. The vertical dotted red line depicst the start of the PAM site with arrow pointing to the direction of insertion for target *eyfp*-2, horizontal red dotted line depicts $1x10^4$ RPM.

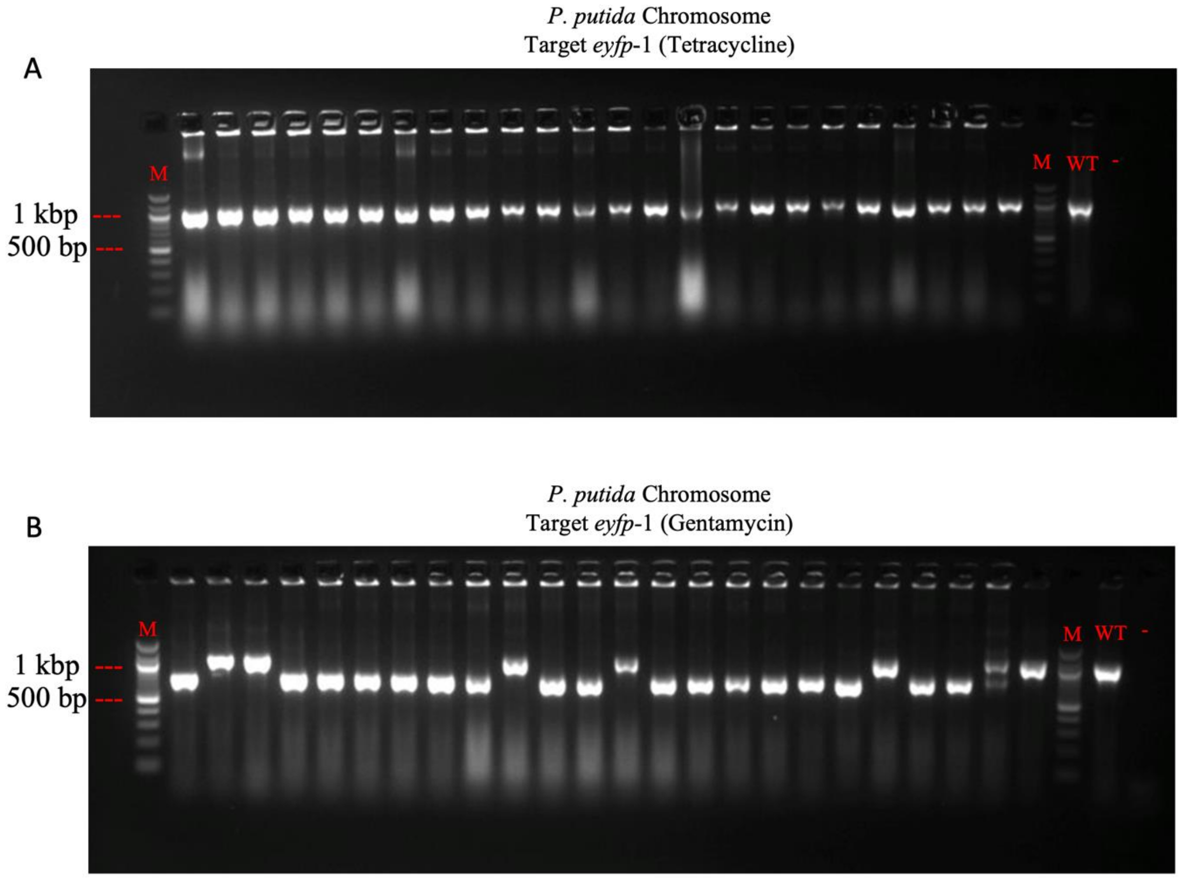


**Supplementary Figure S6. *P. putida* CasTn target *eyfp*-1 genotyping.** *P. putida* CasTn target *eyfp*-1 genotyping. The change in amplicon size from 1.1 Kbp (WT *eyfp)* to ~700 bp shows successful on-target transposition for target *eyfp*-1. Two *eyfp*-1 pDonor plasmids were tested in *P. putida* KT2440 pBLlSh2 carrying a transposon with a tetracycline **(A)** or gentamycin **(B)** selectable marker. PCR amplification was separated on a 2% agarose gel with 100 bp DNA ladder (M), using *P. putida* KT2440 *eyfp* pBLlSh2 (WT) and a no template **(-)** controls.

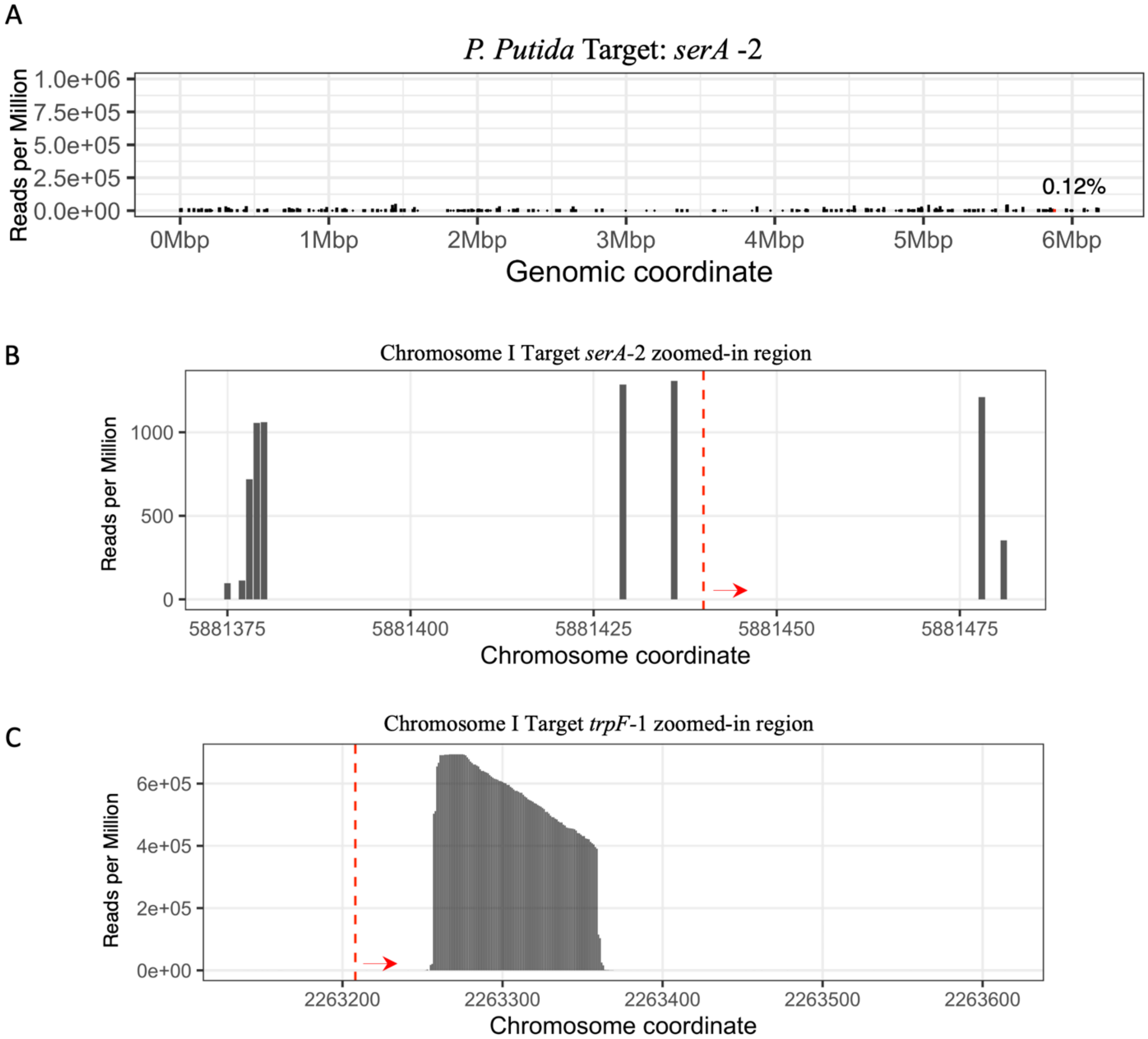


**Supplementary Figure S7. *P. putida* TnSeq results for targets *serA*-2 and *trpF*-1. (A)** TnSeq results for targets *serA*-2 with on-target reads (red bar) and frequency (%) across the whole genome. Zoomed-in view of transposon insertions for target *serA*-2 **(B)** and *trpF*-1 **(C)**; start of gene (red dashed-line) and direction of gene (red arrow) are depicted.

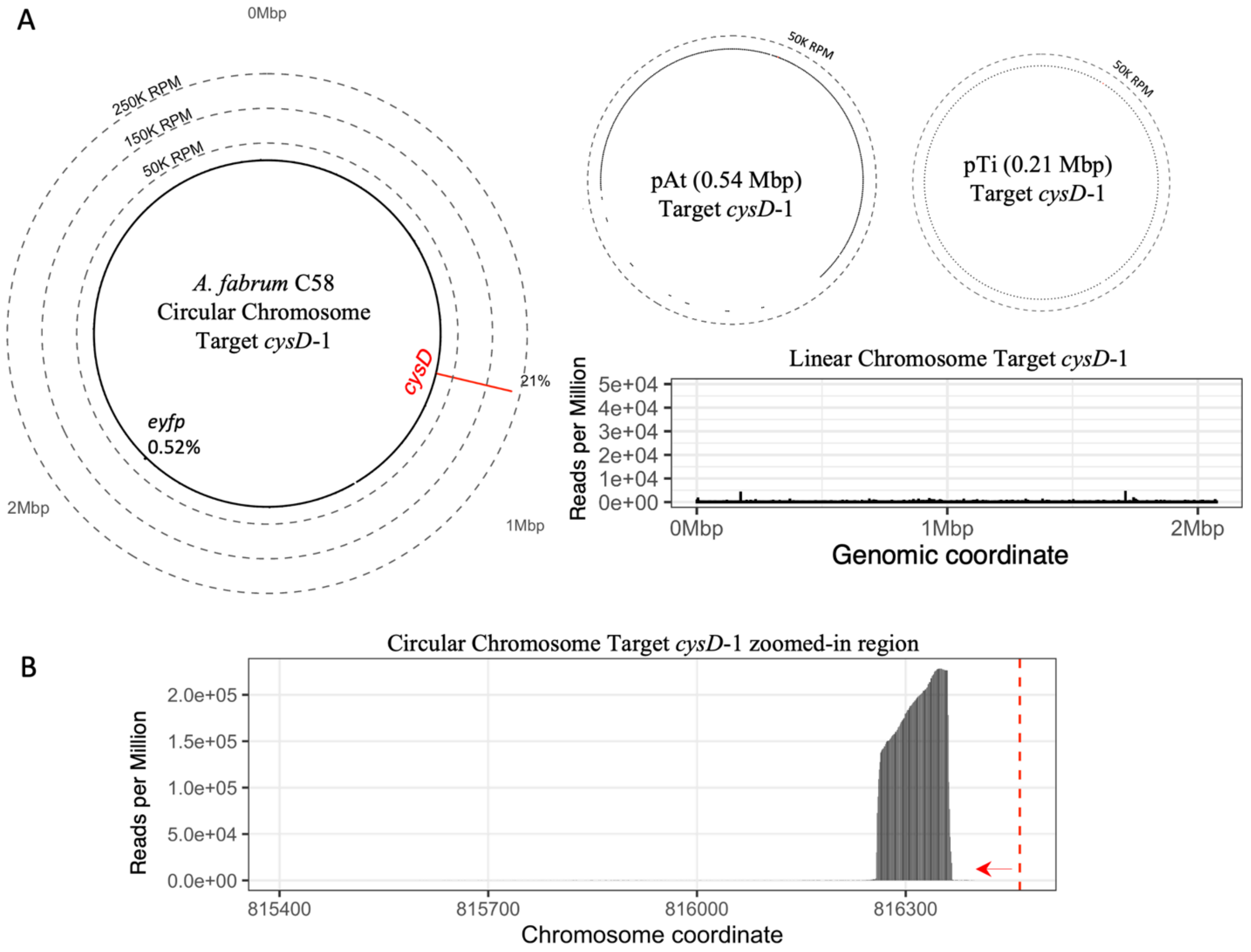


**Supplementary Figure S8.** ***A. fabrum*** **amino acid biosynthesis gene target results.** **(A)** Transposon sequencing results for target *cysD-1* across *A. fabrum* C58 circular chromosome, linear chromosome, and native plasmids pTi and pAt in reads per million (RPM). In the circular chromosome, target *cysD* insertions are depicted by red line accounting for 21% of all reads across the whole genome. **(B)** Zoomed-in view of transposon insertions in *cysD,* the start of the gene (red dashed-line) and direction of the gene (red arrow) are depicted.

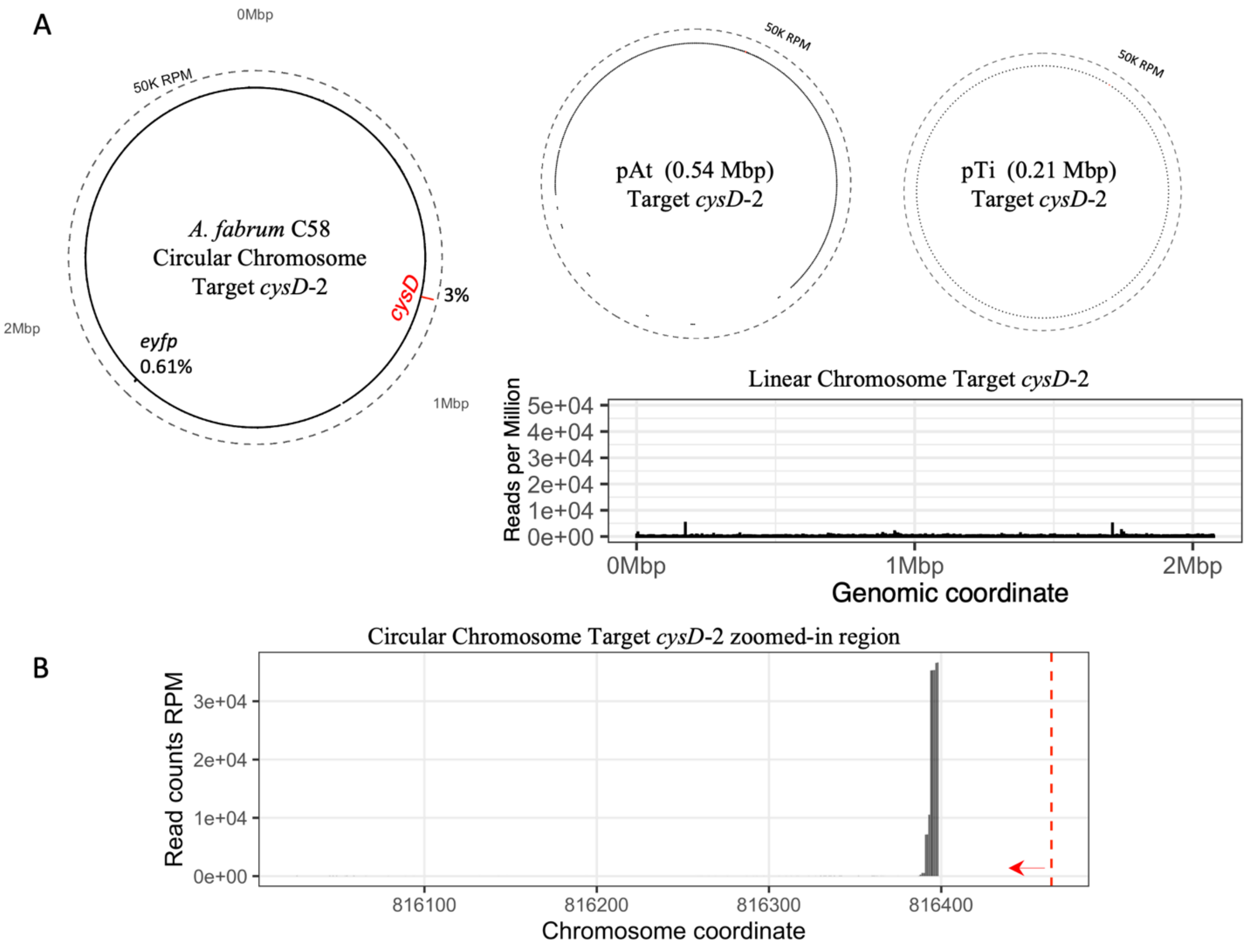


**Supplementary Figure S9. *A. fabrum* TnSeq results for target *cysD*-2. (A)** Transposon sequencing results for target *cysD*-2 across *A.fabrum* C58 circular chromosome, linear chromosome, and native plasmids pTi and pAt in reads per million (RPM). In circular chromosome, target *cysD* insertions are depicted by red line accounting for 3% of all reads in dataset across the whole genome. **(B)** Zoomed-in view of transposon insertions in *cysD,* start of gene (red dashed-line) and direction of gene (red arrow) are depicted.

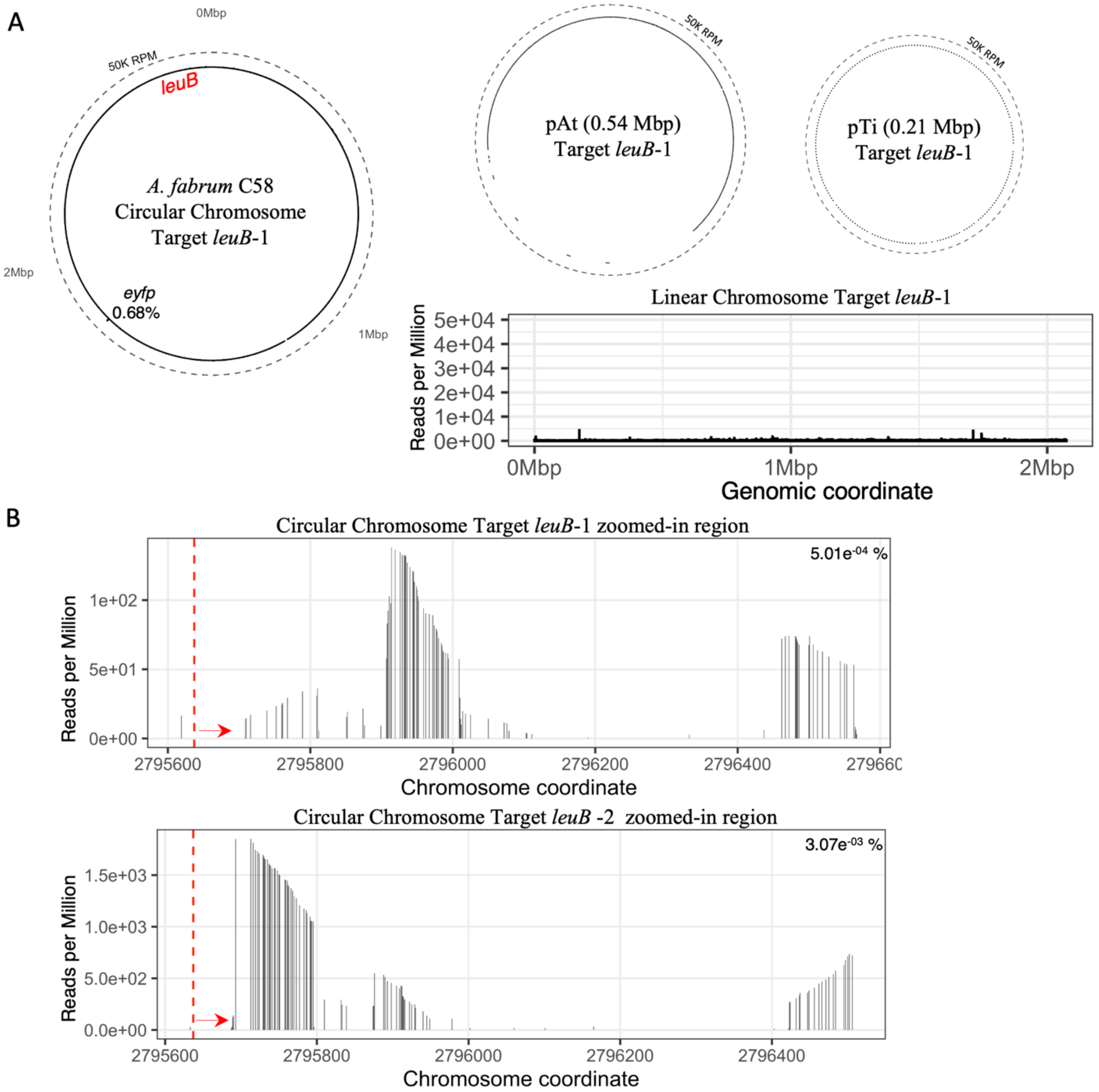


**Supplementary Figure S10. *A. fabrum* TnSeq results for targets *leuB*-1 and *leuB*-2. (A)** Transposon sequencing results for target *lueB*-1 across *A.fabrum* C58 circular chromosome, linear chromosome, and native plasmids pTi and pAt in reads per million (RPM). In the circular chromosome, the region containing the target gene *leuB* is marked (2.7 Mbp). **(B)** Zoomed-in view of transposon insertions in *leuB*-1 and *leuB*-2 start of gene (red dashed-line) and direction of gene (red arrow) are depicted. In the top right corner is the percentage of reads that map to the region shown across the whole genome.

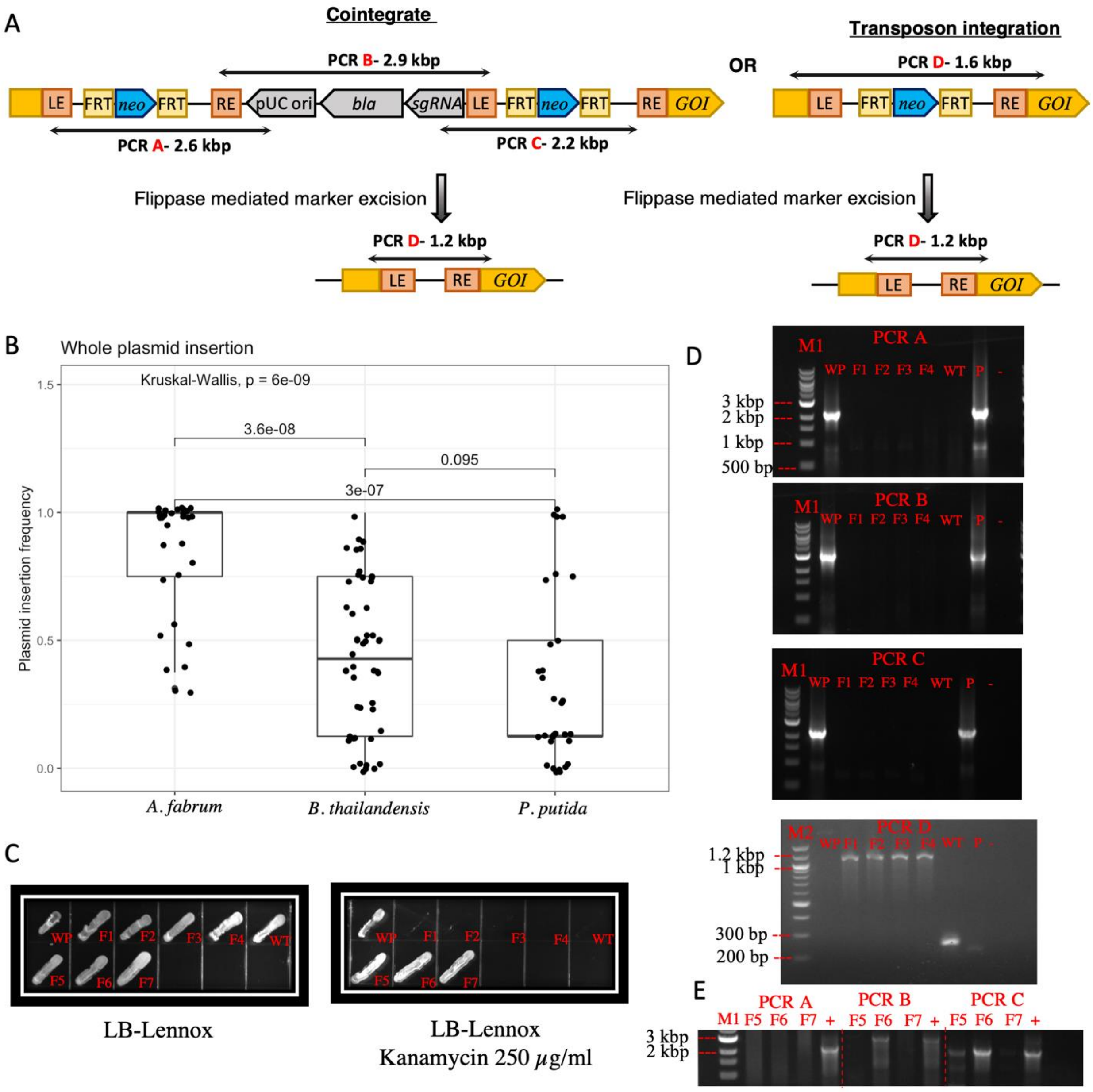


**Supplementary Figure S11. Whole-plasmid integration and Flp mediated excision. (A)** Schematic of whole-plasmid insertion versus transposon integration only. After Flp mediated marker excision, antibiotic-sensitive mutants exhibit loss of whole-plasmid backbone integration. **(B)** Boxplot of whole-plasmid insertion frequency across the three Proteobacteria used in this study. Each dot is a transposon insertion frequency by target in *A. fabrum* (n=11), *B. thailandensis* (n=18), and *P. putida* (n=11). **(C)** Colonies were patched in LB (Left) or LB-Kanamycin (Right) for *B. thailandensis trpA*-1 whole-plasmid integration mutants (WP), Flp mediated mutants (F1-F7) and *B. thailandensis* pKSh5 (WT). **(D)** PCR amplifications of whole-plasmid integration (PCR A-C, 0.8% agarose) and *trpA* gene (PCR D, 2% agarose) with 1 kbp DNA ladder (M1) or 100 bp DNA ladder (M2), using *B. thailandensis* pKSh5 (WT, 216 bp), pUCShDon2 pDonor (P) and no template (-) controls. **(E)** PCR amplifications of whole-plasmid integration (PCR A-C, 0.8% agarose) of incomplete Flp marker excision mutants (F5-F7) with 1 kbp DNA ladder (M1) and pUCShDon2 pDonor (+) control.

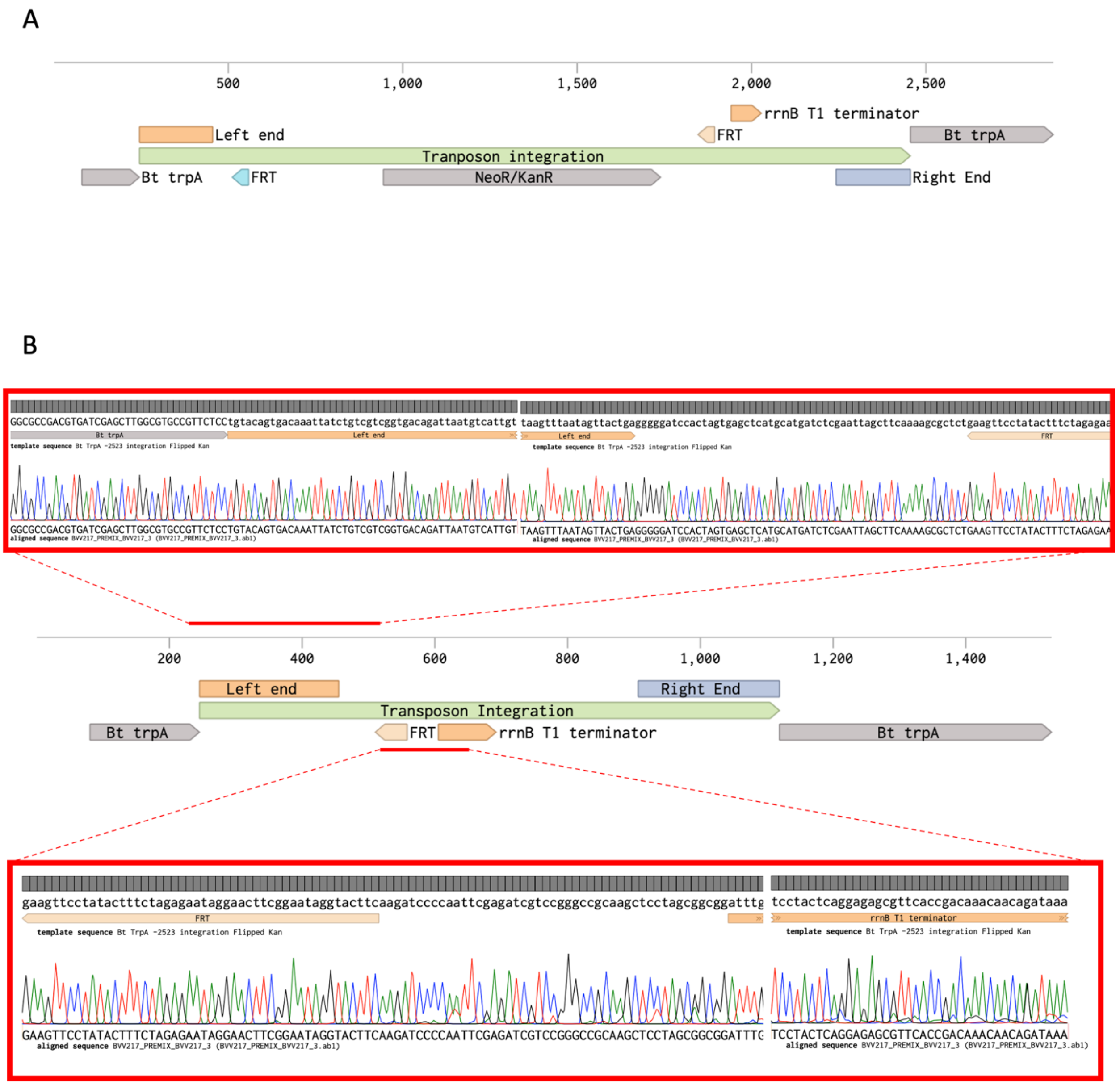


**Supplementary Figure S12. Flp mediated antibiotic marker excision Sanger sequencing. (A)** Schematic of a *B. thailandensis trpA*-1 transposon mutant. FRT (Flp recognition target) sites flank the transposon cargo gene for kanamycin resistance. **(B)** After Flp-mediated marker excision, a *B. thailandensis trpA*-1 kanamycin sensitive mutant was sent for Sanger sequencing. A clean marker excision was observed from the left end of the transposon up to the rrnB1 T1 terminator.

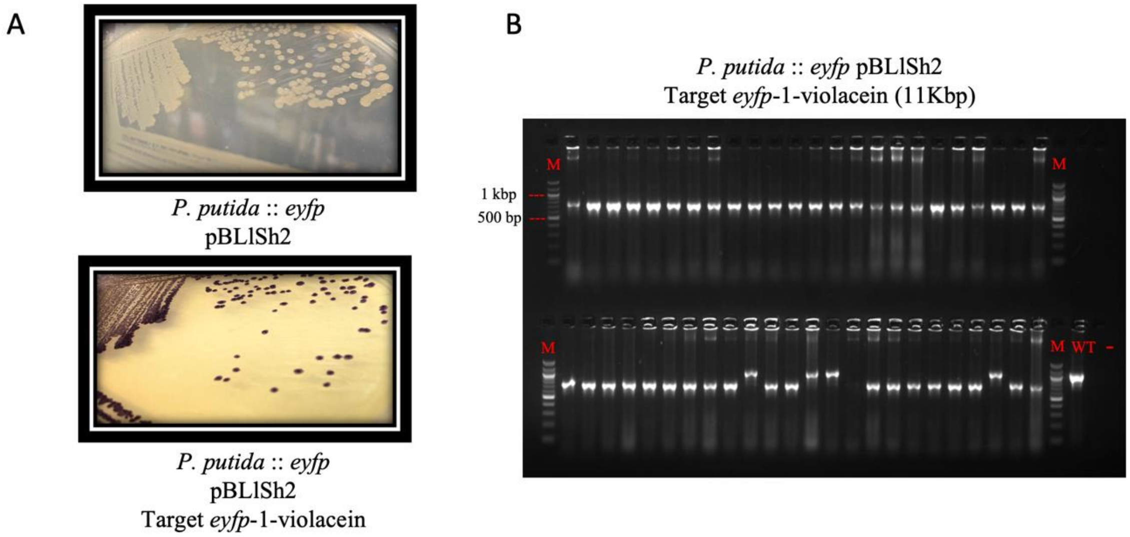


**Supplementary Figure S13. *P. putida* violacein pathway transposon insertion. (A)** *P. putida* pBLlSh2 in LB - Kanamycin (top) and *P. putida* violacein transposon mutant in LB- Gentamycin (bottom) depicting purple colonies, a product of the violacein pathway. **(B)** Target *eyfp*-1 violacein transposon insertion PCR (Supplementary Figure S3) ran on 0.8% agarose gel with 100 bp ladder (M), *P. putida* pBLlSh2 (WT) and no template (-) controls.

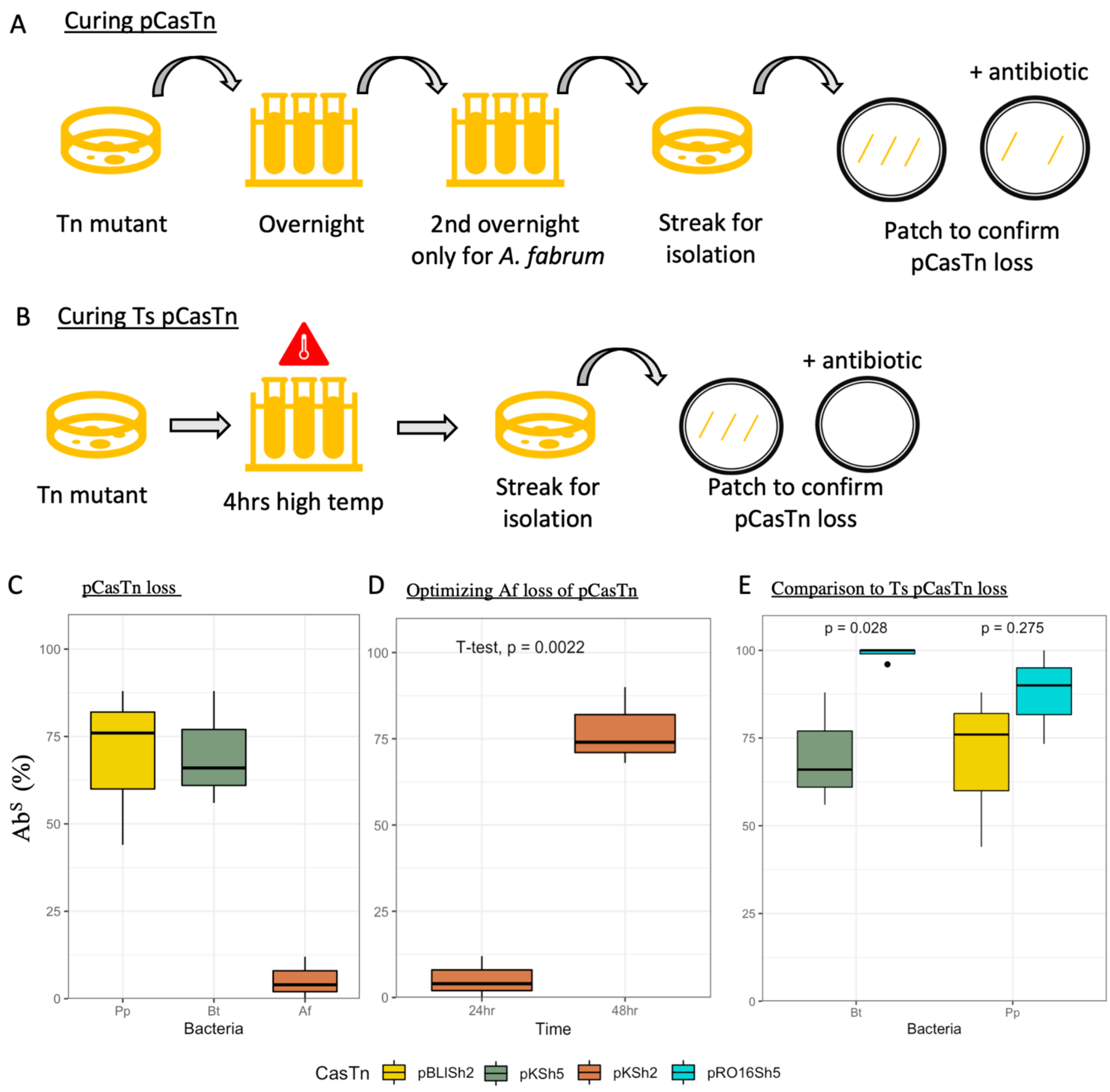


**Supplementary Figure S14. Curability of helper plasmid (pCasTn). (A)** Schematic of pCasTn loss procedure through rich media passage. **(B)** Schematic of Ts pCasTn loss procedure through growth at high temperature. **(C)** Loss of pCasTn after one round of overnight growth, followed by streak for isolation. Isolated colonies were tested for antibiotic sensitivity ($Ab^S$) for the pCasTn marker, results presented as a percentage of the population tested for *Pseudomonas putida* (n=150), *Burkholderia thailandensis* (n=150) and *Agrobacterium fabrum* (n=150). **(D)** Results comparing the first round (24hrs) vs the second round (24hrs) of passage for pCasTn loss in *Agrobacterium fabrum* (n=150). **(E)** Comparison of pCasTn loss after one round of passage (pBLISh2, pKSh5) versus 4hrs of outgrowth at an eleveated temperature (pRO16Sh5) in *Burkholderia thailandensis* (n=100) and *Pseudomonas putida* (n=90).

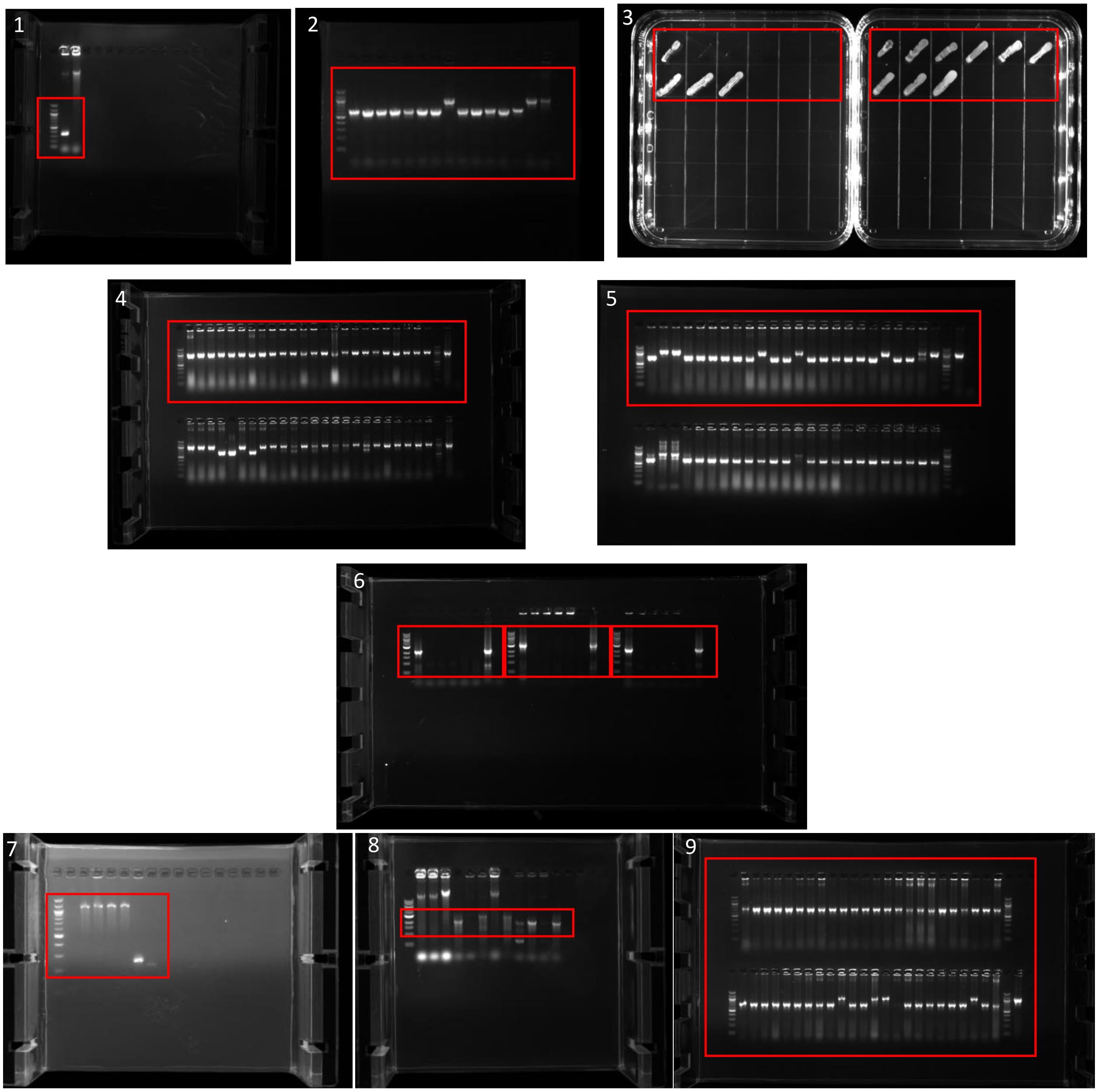


**Supplementary Figure S15. Source data for electrophoresis gels and growth patches in supplementary file.** Picture 1 (Supplementary Figure S2), picture 2 (Supplementary Figure S3), picture 3 (Supplementary Figure S11C), pictures 4-5 (Supplementary Figure S5), pictures 6-8 (Supplementary Figure S11D-E) and picture 9 (Supplementary Figure S12).

| Day | CasTn timeline | HR timeline (example – *sacB* counterselection) |
|---|---|---|
| 1 | 1. Around the horn cloning for **pDonor**: PCR to re-target sgRNA (24 bp), ligate and transformation into *E. coli.* (Parallelizable, High-throughput compatible)<br>2. Electroporate **pCasTn** into Bt. | **Allelic exchange vector cloning:** PCR regions of homology (500bp each) and vector backbone - Assemble fragments - Transform into cloning strain of *E.coli* |
| 2 | Inspect **pDonor** clones for loss of fluorescence..<br>Start overnight culture of *E.coli* with pDonor and Bt + **pCasTn**. | Recover **allelic exchange vector** clones – PCR to check assembly- Start overnight culture of a selected clone. |
| 3 | Miniprep **pDonor** and send for Sanger sequencing-<br>**Integration experiment:** Electroporate Bt + **pCasTn** with ~50 ng pDonor – Plate onto rich media with antibiotic selecting for transposon. | Miniprep **allelic exchange vector** and send for Sanger sequencing. Transform plasmid into *E.coli* with conjugation capability (e.g., RHO3 – DAP auxotroph). |
| 4 | Recover Bt Tn mutants- confirm insertion through PCR. Finished mutant.<br>*Optional: grow culture for marker excision or 2$^{nd}$ integration using CasTn.* | Streak out *E.coli* RHO3 carrying the allelic exchange plasmid in rich media with DAP and antibiotics, incubate for 48hrs. |
| 5 | **Marker excision** of Bt mutant - Transform with flippase (Flp) recombinase carrying plasmid (e.g. pFLpe2[3]). Plate on rich media + Zeo 2,000 and 0.2% L-rhamnose. Incubate 48hrs at 30°C. | Streak out Bt in rich media with DAP, incubate overnight. |
| 6 | Incubation day. | **Conjugation** of Bt and *E.coli* RHO3 + **allelic exchange plasmid**. Incubate 4-6hrs, collect bacteria, spread on plate without DAP but with antibiotic and incubate overnight. |
| 7 | Bt mutant + pFlpe2 colonies appear. Patch into rich media plates with and without the transposon antibiotic. Incubate overnight at 37°C. | **Bt merodiploid** colonies arise – PCR to verify first crossover location on genome is correct. |
| 8 | PCR verify antibiotic sensitive mutants for loss of antibiotic marker.<br>Finished marker excision. | Select sucrose sensitive mutants and plate onto media with sucrose to stimulate second cross-over, incubate for 48 hrs. |
| 9 | | Incubation day. |
| 10 | | Sucrose-resistant colonies are possible double cross-over mutants. Streak and incubate overnight on media without antibiotics. |
| 11 | | Colonies should be patched onto media and check for antibiotic sensitivity, as the marker should be lost on second crossover. |
| 12 | | PCR colonies that show both sucrose resistance and antibiotic sensitivity to verify genotype that could be either wild-type or mutant allele. Finished mutant. |

**Supplementary Figure S16. Timeline of CasTn protocol.** In comparison to homologous recombination (HR) protocol[7,8] in *B. thailandensis*.

| Attributes | CasTn (this paper) | ShCAST[9] | INTEGRATE[10] | MUCICAT[11] |
|---|---|---|---|---|
| sgRNA cloning | around-the-horn PCR | Not described | Golden gate cloning | Golden gate cloning |
| System set up | Two plasmid<br>pHelper: ShCasTn<br>pDonor: sgRNA + transposon (suicide vector) | Two plasmid<br>pHelper: sgRNA + ShCasTn<br>pDonor: transposon | One plasmid (pSPIN)<br>sgRNA + transposon + CasTn<br>Three plasmid<br>pQCascade + sgRNA<br>pTnsABC<br>pDonor (transposon) | Three plasmid<br>pTnsABC<br>pDonor (transposon)<br>pQCascade (sgRNA + Cas + *tniQ*) |
| Direction of insertion (downstream of PAM) | Directionally inserted transposon (left to right end). | Directionally inserted transposon (left to right end). | Random transposon insertion direction. | Directional and random dependent on system used. (INTEGRATE or ShCAST) |
| Max payload size tested | >10Kbp | ~2Kbp | >10Kbp | ~2Kbp |
| Organisms tested | *B. thailandensis*,<br>*A. fabrum*, *P. putida* | *E. coli* | *K. oxytoca*, *P. putida* | *E. coli, Tatumella citrea* |

**Supplementary Figure S17. System comparisons.** Table of comparison of Cas-associated transposase systems.

**Legends Datasets**

**Dataset S1. Raw phenotype and genotype results per mutant.** Raw data of CasTn transposon mutants for growth and PCR reactions. Growth_LB= Growth in LB under antibiotic selection, Growth_M9= Growth in M9, Growth_M9_AA =Growth in M9 + amino acid supplement, (0= No growth, 1=Growth). RE-LE PCR= PCR of genome to LE or RE transposon end in on-target region, WT_PCR= Amplification of unmutated target region, pUC_PCR = PCR for whole pUC plasmid insertion (0= No band, 1= Band present). Fluorescence = Phenotypic assessment of loss of fluorescence (loss of fluorescence = 0, fluorescence =1). LE-eyfp DT= For *Burkholderia thailandensis* multiplex transposition, both *trpA* and *eyfp* RE-LE_PCR were assessed. Results are RE-LE-PCR for *trpA*-1 target and LE-eyfp for *eyfp*-2 target (0= No band, 1= Band present).

**Dataset S2. Plasmid sequences.** Plasmid sequences for donor plasmids with target *eyfp*-1 and CasTn plasmids used in this study. Target *efyp*-1 is emphasized by upper case letters in pDonor sequences. Addgene accession numbers are given.

**Dataset S3. Transformation efficiencies.** Individual transformation efficiencies for each bacterium, CasTn system and target are given in Colony forming units (CFU) per µg of DNA transformed. Bt= *Burkholderia thailandensis,* Pp= *Pseudomonas putida*, Af= *Agrobacterium fabrum.*

**Dataset S4. TnSeq library statistics.** Basic statistics on TnSeq library reads including sample name, bacteria, target, total raw reads, reads after quality filtering and reads mapped to the genome.

**Dataset S5. Genomic locations of targets.** Coordinate positions for all target genes used in this study in *Burkholderia thailandensis, Pseudomonas putida*, and *Agrobacterium fabrum.*

**Dataset S6. Off-target analysis.** Top off-target region for each TnSeq experiment, transposon insertion reads are binned by 1Kbp. Includes off-target % across the whole genome, genomic coordinates and genes found in the off-target region.

**Dataset S7. Insertion distance.** Insertion distance (bp) from 5'PAM3' site to transposon left end insert in genome resolved through Sanger sequencing of individual mutants. Bt= *Burkholderia thailandensis,* Pp= *Pseudomonas putida*, Af= *Agrobacterium fabrum.*

**Dataset S8. Curing pCasTn.** Data for curing pCasTn through passages in media. Grew_LB= Growth in LB, No-Growth-CasTn-Antibiotic = Patch did not grow in media plus antibiotic selecting for pCasTn, Cured = Percent of population cured from pCasTn. Temp = Temperature the culture was grown at (°C), Time = Time in hours grown at temperature (Temp).
